# The first days in the life of naïve human B-lymphocytes infected with Epstein-Barr virus

**DOI:** 10.1101/666297

**Authors:** Dagmar Pich, Paulina Mrozek-Gorska, Mickaël Bouvet, Atsuko Sugimoto, Ezgi Akidil, Adam Grundhoff, Stephan Hamperl, Paul D. Ling, Wolfgang Hammerschmidt

**Affiliations:** Research Unit Gene Vectors, Helmholtz Zentrum München, German Research Center for Environmental Health and German Center for Infection Research (DZIF), Partner site Munich, Marchioninistr. 25, D-81377 Munich, Germany; Heinrich Pette Institute, Leibniz Institute for Experimental Virology, Martinistr. 52, D-20251 Hamburg, Germany; Institute of Epigenetics and Stem Cells, Helmholtz Zentrum München, German Research Center for Environmental Health, Marchioninistr. 25, D-81377 Munich, Germany; Department of Molecular Virology and Microbiology, Baylor College of Medicine, Houston, Texas, United States

## Abstract

Epstein-Barr virus (EBV) infects and activates resting human B-lymphocytes, reprograms them, induces their proliferation, and establishes a latent infection in them. In established EBV-infected cell lines many viral latent genes are expressed. Their roles in supporting the continuous proliferation of EBV-infected B cells *in vitro* are known, but their functions in the early, pre-latent phase of infection have not been investigated systematically. In studies during the first eight days of infection using derivatives of EBV with mutations in single genes of EBVs we found only EBNA2 to be essential for activating naïve human B-lymphocytes, inducing their growth in cell volume, driving them into rapid cell divisions, and preventing cell death in a subset of infected cells. EBNA-LP, LMP2A and the viral microRNAs have supportive, auxiliary functions, but mutants of LMP1, EBNA3A, EBNA3C, and the noncoding EBER RNAs had no discernable phenotype compared with wild-type EBV. B cells infected with a double mutant of EBNA3A and 3C had an unexpected proliferative advantage and did not regulate the DNA damage response (DDR) of the infected host cell in the pre-latent phase. Even EBNA1 which has very critical long-term functions in maintaining and replicating the viral genomic DNA in established cell lines, was dispensable for the early activation of infected cells. Our findings document that the virus dose is a critical parameter and indicate that EBNA2 governs the infected cells initially and implements a strictly controlled temporal program independent of other viral latent genes. It thus appears that EBNA2 is sufficient to control all requirements for clonal cellular expansion and to reprogram human B-lymphocytes from energetically quiescent to activated cells.

**Author summary:** The preferred target of Epstein-Barr virus (EBV) are human resting B-lymphocytes. We found that their infection induces a well-coordinated, time-driven program that starts with a substantial increase in cell volume followed by cellular DNA synthesis after three days and subsequent rapid rounds of cell divisions on the next day accompanied by some DNA replication stress (DRS). Two to three days later the cells decelerate and turn into stably proliferating lymphoblast cell lines. With the aid of 16 different recombinant EBV strains we investigated the individual contributions of EBV’s multiple latent genes during early B-cell infection and found that many do not exert a detectable phenotype or contribute little to EBV’s pre-latent phase. The exception is EBNA2 that is essential in governing all aspects of B-cell reprogramming. EBV relies on EBNA2 to turn the infected B-lymphocytes into proliferating lymphoblasts preparing the infected host cell for the ensuing stable, latent phase of viral infection. In the early steps of B-cell reprogramming viral latent genes other than EBNA2 are dispensable but some, EBNA-LP for example, support the viral program and presumably stabilize the infected cells once viral latency is established.

## Introduction

In 1964, Epstein, Achong, and Barr identified a new herpes virus in a cell line derived from Burkitt’s lymphoma (Epstein et al., 1964) and three years later two groups independently found that this virus, now termed Epstein-Barr virus (EBV), can transform human primary B-lymphocytes into lymphoblastoid cell lines (LCLs) that proliferate in culture (Henle et al., 1967; Pope, 1967; Pope et al., 1968).

Initially, many groups have focused on the identification of the viral genes and factors expressed in the latently infected and proliferating human B cells, i.e. EBV’s latent genes. This explorative period was followed by a second wave of publications starting in 1989 (Hammerschmidt and Sugden, 1989; Cohen et al., 1989), which studied the genetic requirements of the identified latent viral genes in this B cell model. We found that EBNA2 was an essential viral gene (Kempkes et al., 1995c) and other early publications suggested that the latent membrane protein 1 (LMP1), Epstein-Barr nuclear antigen 3A (EBNA3A) and EBNA3C were essential for the generation of LCLs from *in vitro* infected human B-lymphocytes, whereas LMP2A, LMP2B, EBNA3B, and the two non-coding RNAs, EBER1 and 2, were dispensable but contributed to this process (Kieff and Rickinson, 2007). With the emergence of more sophisticated genetic approaches it was possible to study EBV’s latent genes more accurately. The synthetic assembly of mini-EBV genomes (Kempkes et al., 1995b; Kempkes et al., 1995a; Kilger et al., 1998) and the eventual cloning of the entire EBV genome in E. coli (Delecluse et al., 1998) opened the field for the unlimited genetic analysis of all viral genes (and EBV’s cis-acting elements) (Feederle et al., 2010). From these studies (and in the context of an otherwise genetically unaltered EBV genome) it became clear that LMP1 (Dirmeier et al., 2003), EBNA3A (Kempkes et al., 1995b; Hertle et al., 2009), but also EBNA3C in B-cells from p16 deficient donors (Skalska et al., 2010) are dispensable and only EBNA2 is absolutely essential to induce and maintain LCLs from *in vitro* EBV-infected B-lymphocytes (Altmann and Hammerschmidt, 2005).

Very much in contrast to many viruses EBV does not start the *de novo* synthesis of virus progeny upon infection, rather, it initiates a latent infection (Kalla et al., 2010; Kalla et al., 2012). It was therefore surprising to learn that certain lytic viral genes are expressed in newly infected B cells (Wen et al., 2007; Kalla et al., 2010; Jochum et al., 2012b; Jochum et al., 2012a). Their expression is transient, only, but some are essential for the emergence of lymphoblastoid cell lines (Altmann and Hammerschmidt, 2005) suggesting that EBV uses a set of viral genes in the first few days of infection that differs from the set of viral genes expressed in established, stable and latently infected lymphoblastoid cell lines (Hammerschmidt, 2015).

Here, we sought to analyze the individual genetic contributions of all latent viral genes to the activation and transformation of human B-lymphocytes during the first eight days following infection. Our experiments suggest that the reprogramming of newly infected cells, which follows a very strict, time-controlled scheme is orchestrated solely by EBNA2, a viral factor that has been shown to be essential for maintaining B cell transformation, previously (Kempkes et al., 1995c).

## Results

### The virus dose is an important parameter of cell survival and proliferation in the pre-latent phase of EBV infection

We started this study by refining our previous data (Steinbrück et al., 2015) to determine the parameters of infection and to identify the optimal ratio of EBV versus human primary B-lymphocytes. We infected them with different multiplicities of infection (MOIs) of wt/B95.8 (2089) EBV (Tab. 1) and counted the absolute number of viable lymphocytes and emerging B blasts by FACS employing calibrated APC beads (Calibrite, Becton Dickinson) as a volume standard. In this analysis we also investigated the binding of Annexin V, an early indicator of apoptosis. Concomitantly, we determined the fraction of EBNA2-positive B cells. Towards this end we fixed and permeabilized the cells, stained them with an anti-EBNA2 antibody coupled with Alexa647 and analyzed the stained cells by FACS.

**Table 1.** Features of the EBV strains used in infection experiments.

| EBV strain | Genotype | Specification | Genealogy | Reference |
| --- | --- | --- | --- | --- |
| wt/B95.8 (2089) | wild-type, 13 miRNAs, *hpt* | reference EBV strain |  | (Delecluse et al., 1998) |
| wt/B95.8 (6001) | wild-type, 13 miRNAs, *pac* | reference EBV strain | based on wt/B95.8 (2089) | this study |
| r_wt/B95.8 (6008) | wild-type, 44 miRNAs, *pac* | deletion in wt/B95.8 (2089) restored | based on wt/B95.8 (2089) | this study |
| wt/B95.8 (5750) | wild-type, 13 miRNAs, *hpt* | six BamHI-W-repeats | based on wt/B95.8 (2089) | this study |
| ΔEBNA-LP (5969) | EBNA-LP knockout, *pac* | six BamHI-W-repeats, in-frame translational stop codon in each W2 exon of EBNA-LP | based on wt/B95.8 (2089) | this study |
| ΔEBNA2 (5968) | EBNA2 knockout, *pac* | AUG to TGA mutation of EBNA2's translational start codon | based on wt/B95.8 (2089) | this study |
| ΔmiR (4027) | no viral miRNAs, *hpt* | scrambled pre-miRNA loci | based on wt/B95.8 (2089) | (Seto et al., 2010) |
| r_ΔmiR (6338) | no viral miRNAs, *pac* | scrambled pre-miRNA loci | based on r_wt/B95.8 (6008) | this study |
| ΔLMP1 (2597) | LMP1 knockout, *hpt* | deletion of *BNLF1* | based on wt/B95.8 (2089) | (Dirmeier et al., 2003) |
| ΔLMP2A (2525) | LMP2 knockout, *hpt* | Cre-mediated deletion of first exon of LMP2A | based on wt/B95.8 (2089) | (Mancao et al., 2005) |
| ΔEBER (6431) | knockout of EBER1 and 2, *pac* | insertional mutagenesis | based on r_wt/B95.8 (6008) | this study |
| ΔEBER/ΔmiR (6432) | knockout of EBER1,2, miRNAs, *pac* | scrambled pre-miRNA loci, insertional mutagenesis | based on r_ΔmiR (6338) | this study |
| ΔEBNA3A (6077) | knockout of EBNA3A, *pac* | insertional mutagenesis of EBNA3C | based on wt/B95.8 (6001) | this study |
| ΔEBNA3C (6123) | knockout of EBNA3C, *pac* | two in-frame stop codons in exon 1 and 3 of EBNA3C | based on wt/B95.8 (6001) | this study |
| ΔEBNA3A/C (6331) | knockout of EBNA3A and 3C, *pac* | insertional mutagenesis of EBNA3A | based on ΔEBNA3C (6123) | this study |
| ΔEBNA1 (6285) | knockout of EBNA1, *pac* | AUG to TAG mutation of EBNA1's translational start codon | based on r_wt/B95.8 (6008) | this study |

Delecluse HJ, Hilsendegen T, Pich D, Zeidler R, Hammerschmidt W (1998) Propagation and recovery of intact, infectious Epstein-Barr virus from prokaryotic to human cells. *Proc Natl Acad Sci U S A* 95:8245–8250
Dirmeier U, Neuhiel B, Kilger E, Reisbach G, Sandberg ML, Hammerschmidt W (2003) Latent membrane protein 1 is critical for efficient growth transformation of human B cells by Epstein-Barr virus. *Cancer Res* 63:2982–2989
Mancao C, Altmann M, Jungnickel B, Hammerschmidt W (2005) Rescue of “crippled” germinal center B cells from apoptosis by Epstein-Barr virus. *Blood* 106:4339–4344
Seto E, Moosmann A, Gromminger S, Walz N, Grundhoff A, Hammerschmidt W (2010) Micro RNAs of Epstein-Barr virus promote cell cycle progression and prevent apoptosis of primary human B cells. *PLoS Pathog* 6:e1001063

The concentration of infectious particles in stocks of EBV is measured by infecting Raji cells and determining the fraction of GFP-positive cells by FACS three days later as described in detail in Fig. 2 of our previous work (Steinbrück et al., 2015). In this paper, we also noticed that Daudi cells are much more permissive for EBV infection than Raji cells documenting that we underestimate the virus concentration by at least a factor of ten or more when using Raji cells for virus quantification (Fig. 2B in Steinbrück et al., 2015). In two independent experiments with sorted naïve B-lymphocytes from adenoids we confirmed that an MOI of 0.1 based ‘on green Raji units’ (GRUs) was optimal (Fig. 1A), because this dose reproducibly yielded the highest numbers of B cells eight days post infection (p.i.). Lower MOIs were inferior, but it was unexpected to learn that virus doses beyond 0.1 also resulted in lower cell numbers on day 8 p.i., although the fraction of EBNA2 cells was initially higher compared with cells infected with an MOI of 0.1 (Fig. 1C). Also, Annexin V binding was dramatically increased when the cells were infected with a high MOI of 3.0 (Fig. 1B). We learnt from these experiments that an MOI of 0.1 on Raji cells equates to an MOI of 1.0 or higher on B-lymphocytes. Our results also indicated that EBV’s success is dose-dependent and has a well-defined, but relatively narrow virus-host cell ratio that optimally supports the early survival, activation, and reprogramming of EBV’s target B-lymphocytes. Consequently, we chose an MOI of 0.1 GRU for most subsequent experiments.

**Fig. 1.**
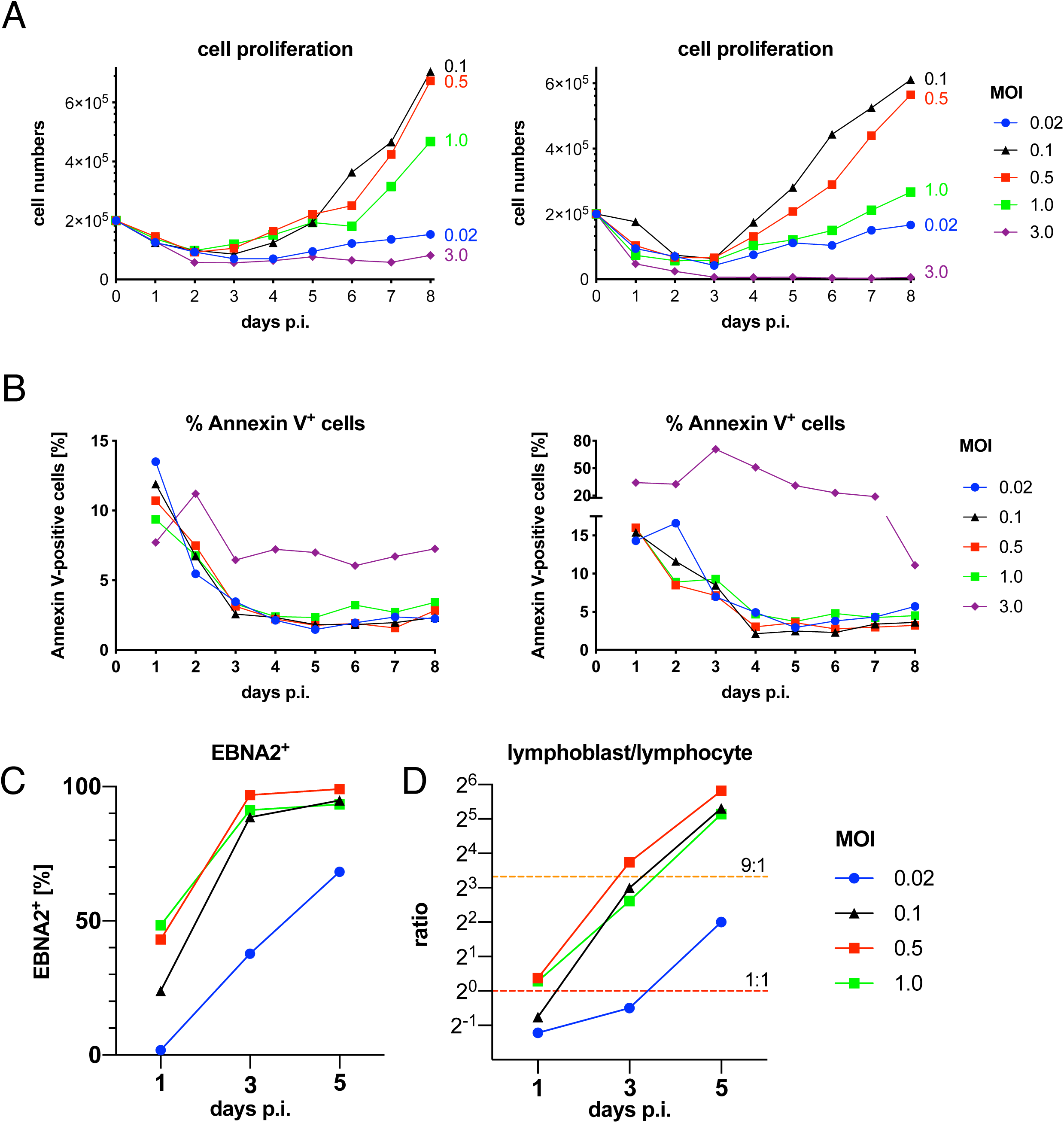
Evaluation of parameters and conditions supporting EBV infection of naive B-lymphocytes. **(A)** Primary naïve B-lymphocytes were sorted and infected with wt/B95.8 (2089) EBV using the indicated multiplicities of infection (MOI). The number of proliferating, growth transformed B cells was recorded by FACS daily as described in Materials and Methods and in Fig. 2A of our recent publication (Steinbrück et al., 2015). Two experiments with B cells from two different donors are shown. **(B)** Annexin V binding of infected B-lymphocytes from the two donors analyzed in panel A is provided. **(C)** The fraction of EBNA2-positive cells in B cells on day 1, 3, and 5 p.i. is shown as a function of MOI. **(D)** The ratio of lymphoblasts versus lymphocytes as determined by forward and sideward scatter FACS analysis was calculated with B cells infected with wt/B95.8 (2089) and different MOIs as indicated on day 1, 3, and 5 p.i.. The horizontal dashed red line indicates a 1 to 1 ratio, the orange line indicates a 9 to 1 ratio of lymphoblast versus lymphocytes. Panels A and B show two representative experiment out of three, panels C and D show one representative experiment out of three.

**Fig. 2.**
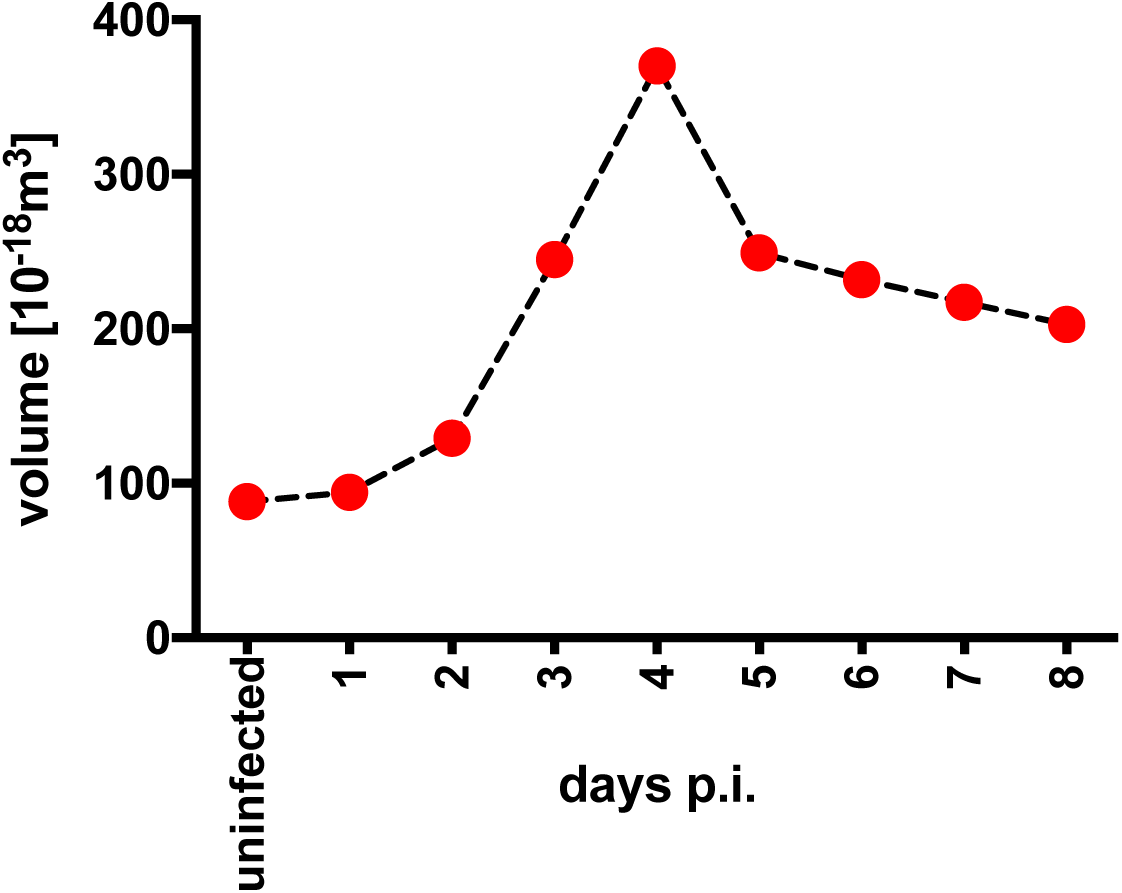
Cell diameter and cell volume of naïve B-lymphocytes infected with EBV. FACS-sorted naïve B-lymphocytes from adenoid tissue were left uninfected or were infected with wt/B95.8 (2089) EBV with an MOI of 0.1 and cultivated for the indicated days. After Ficoll gradient centrifugation, microscopic images of the samples were recorded with a Tali Image-Based Cytometer (Thermo Fisher), analyzed for their cellularity and the cells’ diameters were determined with the aid of suitable calibration beads as described in Materials and Methods. Based on the cell diameter the mean volume of at least 200 cells per time point was calculated assuming perfect spheres.

### EBV reprograms naïve human B-lymphocytes in discrete steps

We isolated primary human B-lymphocytes from adenoid tissue and sorted the fraction of quiescent naïve B cells (IgD^+^/IgH^+^, CD38^-^, CD27^-^) for our infection experiments. The naïve B cells were infected with the recombinant EBV based on the B95.8 strain termed wt/B95.8 (2089) (Delecluse et al., 1998) (Tab. 1) using an MOI of 0.1. We recorded the diameter of the viable cells daily by microscopic imaging and calculated their volume assuming a spherical shape. Uninfected cells had a diameter of 5.5±0.5μm (mean and standard deviation), which gradually increased to close to 8.9μm on day 4 post infection (p.i.), but decreased to 7.5±1.2μm on day 8 p.i. (Fig. 2 and Mrozek-Gorska et al., 2019). Similarly, the inferred volume of the cells increased from 87×10^−18^m^3^ to 357×10^−18^m^3^ on day 4 p.i. and declined to about 220×10^−18^m^3^ (mean values) on day 8 p.i. (Fig. 2) indicating a roughly fourfold change within this period of infection.

We repeated the experiments with sorted naïve B-lymphocytes that were labeled with an intracellular dye (cell trace violet, CTV) prior to infection to monitor the cell doublings over time. Four different EBV strains were used for infection, which represent four versions of the wt/B95.8 strain (Delecluse et al., 1998), but differ in certain genotypic features (Tab. 1). Briefly, wt/B95.8 (2089) and wt/B95.8 (6001) are identical recombinant EBVs but encode *hpt* or *pac* in the virus producer cells conferring resistance to hygromycin B and puromycin, respectively. Based on wt/B95.8 (2089), wt/B95.8 (5750) was engineered to contain six complete copies of the BamHI-W-repeat cluster (see below for details).

The forth versions of the wt/B95.8 strain termed r_wt/B95.8 (6008) adds to our collection of available EBV strains (Tab. 1). The B95.8 reference strain of EBV originates from a lymphoblastoid cell line obtained by infecting marmoset monkey peripheral blood leukocytes with EBV from a patient with infectious mononucleosis (Miller et al., 1972; Miller and Lipman, 1973). The B95.8 EBV strain readily immortalizes human B-lymphocytes and has been studied for decades because the size of its genome was recognized to be smaller than most other EBV field strains, an advantage in the early search for EBV’s immortalizing functions (Baer et al., 1984; Palser et al., 2015). Its genome, however, suffers from a unique deletion (Bornkamm et al., 1980; Heller et al., 1981) that affects the cluster of EBV’s miRNAs (Pfeffer et al., 2004), several protein-encoding viral genes (Parker et al., 1990) and a second copy of EBV’s lytic origin of DNA replication (Hammerschmidt and Sugden, 1988). It has remained uncertain whether these viral functions might contribute to B cell immortalization or establishment of latency. To address this issue we engineered a derivative of wt/B95.8 (2089) termed r_wt/B95.8 (6008) EBV, in which the 12 kb deletion present in the B95.8 EBV strain was restored with the autologous sequences of the M-ABA EBV isolate (Bornkamm et al., 1980) to represent the genetic content of common field strains of EBV (Palser et al., 2015; Farrell, 2015). The DNA sequence of the r_wt/B95.8 (6008) EBV strains corresponds to the reference EBV genomic sequences AJ507799 and NC_007605.

With the four related B95.8 EBV derivates wt/B95.8 (2089), wt/B95.8 (6001), wt/B95.8 (5750), and r_wt/B95.8 (6008) summarized in Table 1, we infected primary sorted naïve B-lymphocytes and analyzed them by FACS daily to record cell doublings, Annexin V binding, and numbers of intact cells for eight days. Cellular DNA synthesis and cell cycle distribution were investigated after metabolic labeling with 5’-bromo-2’-deoxyuridine (BrdU) for one hour and subsequent immunodetection and FACS analysis of the cellular DNA content.

The naïve B-lymphocytes did not divide within the first three days of infection (Fig. 3A) indicating that the increase in cell volume (Fig. 2) reflects the metabolic growth of the infected and activated cells, only. The first cell division occurred on day 4 followed by a short period of rapidly dividing cells until day 6 post infection (p.i.). Thereafter, the dividing cells decelerated considerably to adopt a rate of cell divisions also seen in established lymphoblastoid cell lines. BrdU incorporation was first detected on day 3 p.i. but never earlier, followed by a rapid increase of the fraction of S-phase cells that reached a maximum on day 5 or 6 p.i. (Fig. 3B). Cell numbers dropped dramatically and, in most experiments, up to 70% of the initially infected cells were lost within the first three days of cell culture (Fig. 3C). Cell numbers increased from day 4 onwards in parallel with the onset of cell divisions (Fig. 3A). A high and often variable fraction of cells bound Annexin V within the first three days of cell culture, but starting on day 4 generally more than 90% of the intact viable cells became Annexin V-negative (Fig. 3D). The phenotypes of the cells infected with the four individual wild-type EBV stocks did not differ much, but merely reflected experimental variability of the assay. The novel EBV strain r_wt/B95.8 (6008), which gets as close as possible to an EBV field strain while preserving the context of the prototypic B95.8 EBV does not reveal a discrete phenotype in this set of experiments, but opens new possibilities for investigating EBV-encoded miRNAs.

**Fig. 3.**
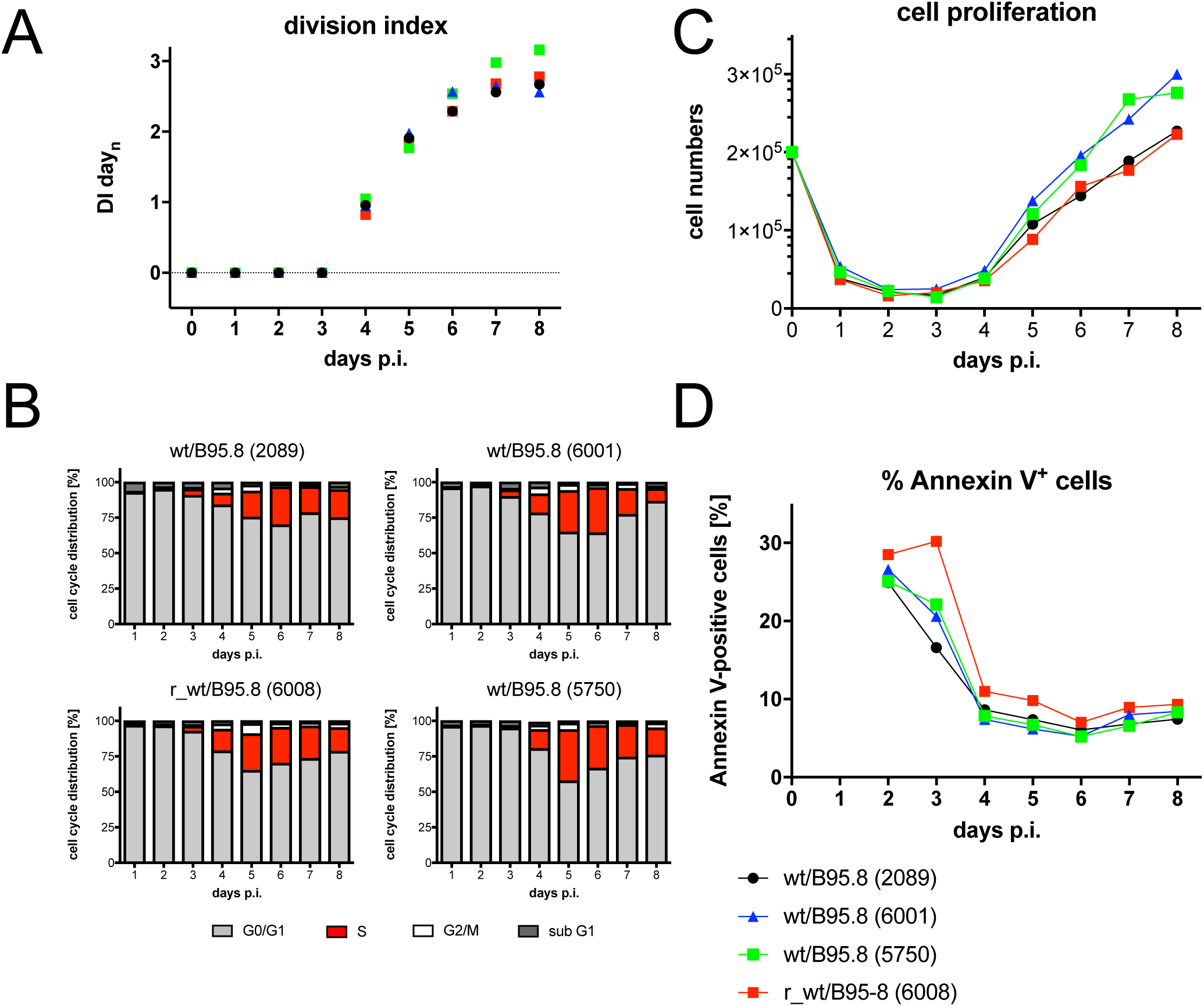
Activation kinetics of naïve B-lymphocytes infected with four different recombinant wild-type EBV strains. Sorted naïve B-lymphocytes isolated from adenoid tissue were loaded with an intracellular dye (cell trace violet, CTV, Thermo Fisher Scientific). The cells were infected with the four indicated EBV strains with an MOI of 0.1. The strains are wild-type with respect to EBV’s latent genes, but have different genotypes as specified in Table 1. **(A)** The kinetics of cell division of the infected cells were analyzed by FACS, the resulting Division Index (DI) was calculated and plotted. DI indicates the average number of cell divisions a cell in the starting populations has undergone including the peak of undivided cells. **(B)** Cells were incubated with 5-Bromo-2’-deoxyuridine (BrdU) for 1 hour prior to harvest and analyzed by FACS after staining with a BrdU specific antibody. The percentage of cells in the different phases of the cell cycle were calculated. **(C)** Cell numbers of viable cells were analyzed by FACS as described (Steinbrück et al., 2015) and plotted. Initial cell numbers in this experiment were 2×105 per well as indicated. **(D)** Annexin V binding of infected cells was analyzed by FACS. One representative experiment out of three experiments with B-lymphocytes from three individual donors is shown.

### Reprogramming of resting B-lymphocytes from peripheral blood

We asked if the phenotype of the EBV-infected naïve B-lymphocytes might reflect their origin from secondary lymphatic tissue. Therefore, we repeated the previous experiment with B cells isolated from peripheral blood, which contains resting, non-activated naïve and memory B-lymphocytes, only. Upon infection of peripheral B-lymphocytes from two donors with wt/B95.8 (2089), we recorded cell size and granularity, cell numbers, division index, and rate of apoptosis by FACS (Fig. S1), but were unable to analyze cellular DNA synthesis due to low initial cell numbers. Similar to sorted naïve B-lymphocytes from adenoid tissue, the majority of peripheral B cells died within the first three days, but the surviving cells were rapidly activated (Fig. S1), started to proliferate on day 4 p.i. and reduced the rate of proliferation on day 6. In general, the percentage of Annexin V-positive cells early after infection was not as high as in naïve B-lymphocytes from adenoids, but the phenotypic differences of infected naïve B-lymphocytes from adenoids versus B-lymphocytes from peripheral blood were minor indicating that B-lymphocytes from different sources apparently followed the identical time scheme.

### CD40 activation and IL-4 induce a comparable program of B cell activation and proliferation

Stimulation of human B-lymphocytes with CD40 ligand (CD40L) and the cytokine IL-4 induces cell activation and unlimited B cell proliferation independent of EBV infection (Banchereau et al., 1991; Wiesner et al., 2008). We reanalyzed this model and found that the kinetics of naïve B cell survival, activation, and proliferation after CD40L and IL-4 stimulation did not differ in principle from EBV-infected resting B-lymphocytes in the first week (Fig. 4). In the course of three independent experiments the fraction of B cells undergoing DNA synthesis peaked on day 4 p.i., but the rate of B cell proliferation was reduced in this model, which resulted in a slow increase of total cell numbers in the observation period (Fig. 4). Again, the onset of cellular DNA synthesis took place on day 3 p.i. and the first cell divisions became apparent one day later.

**Fig. 4.**
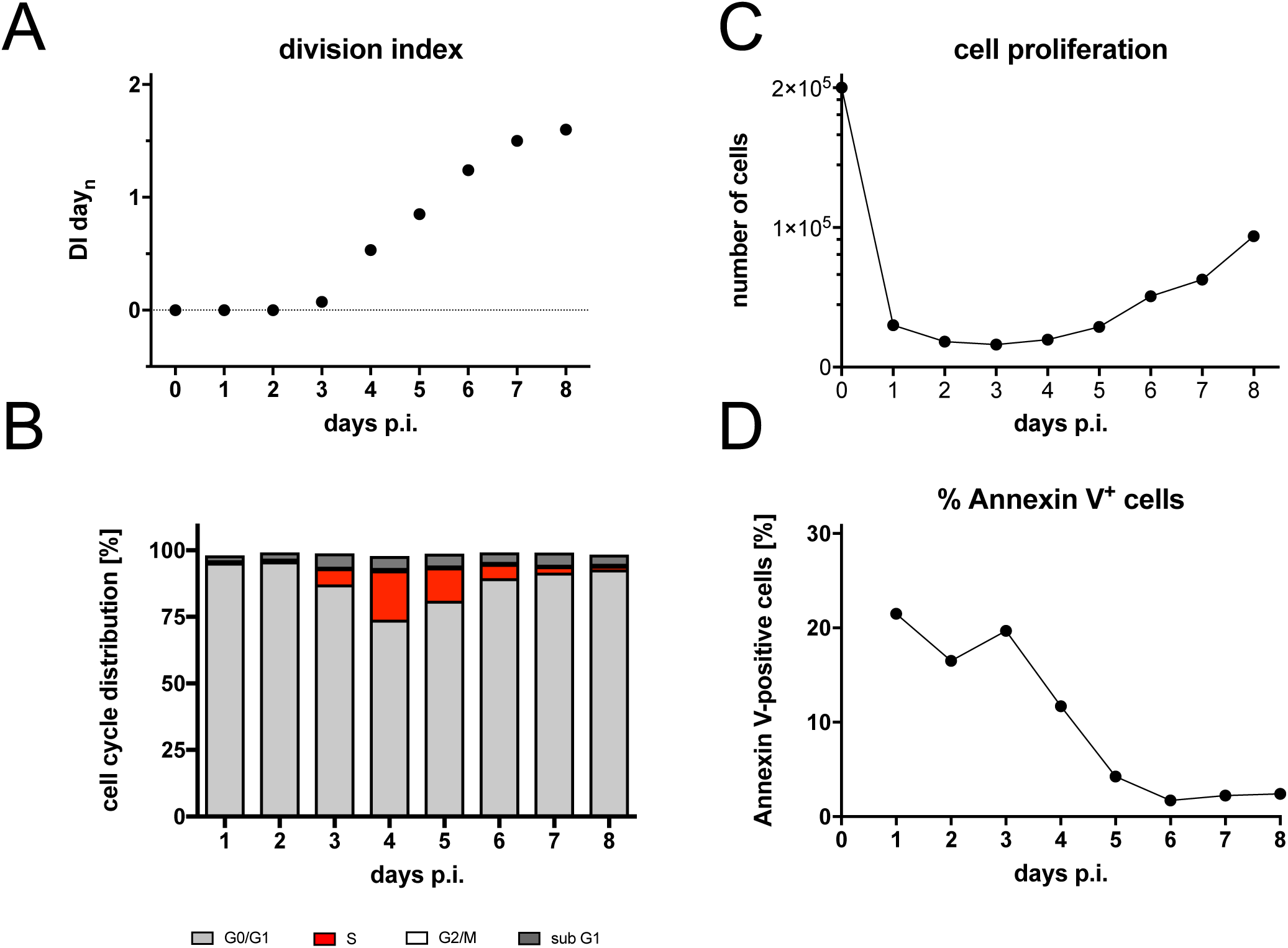
Activation kinetics of naïve B-lymphocytes cultivated on CD40L feeder cells in the presence of IL-4. As in Fig. 3, naïve B-lymphocytes isolated from adenoid tissue were loaded with an intracellular dye and cultivated on CD40L feeder cells with IL-4 as described (Wiesner et al., 2008). **(A)** Division Index is shown. **(B)** Cell cycle distributions of cultivated cells are provided. **(C)** Viable cells were counted by FACS. **(D)** Binding of Annexin V was analyzed by FACS. Shown is one representative experiment out of three with B-lymphocytes from three different B cell donors.

### EBNA2 is essential for entry into reprogramming

EBNA2 has been identified to be essential for B cell immortalization by EBV (Hammerschmidt and Sugden, 1989; Cohen et al., 1989). EBNA2’s functions are also required immediately after EBV infection of B-lymphocytes (Altmann and Hammerschmidt, 2005) and for the continuous propagation of lymphoblastoid cell lines (Kempkes et al., 1995c). We revisited EBNA2’s role in the first week of infection with an EBV mutant incapable of translating EBNA2 mRNA, because it carries a mutation of EBNA2’s translational start codon (Tab. 1). As expected, naïve B-lymphocytes infected with ΔEBNA2 (5968) were not activated and did not survive until day 3 p.i. to synthesize cellular DNA (Fig. 5). The data confirm that EBNA2 is central in all early events in B cell activation and survival ex vivo.

**Fig. 5.**
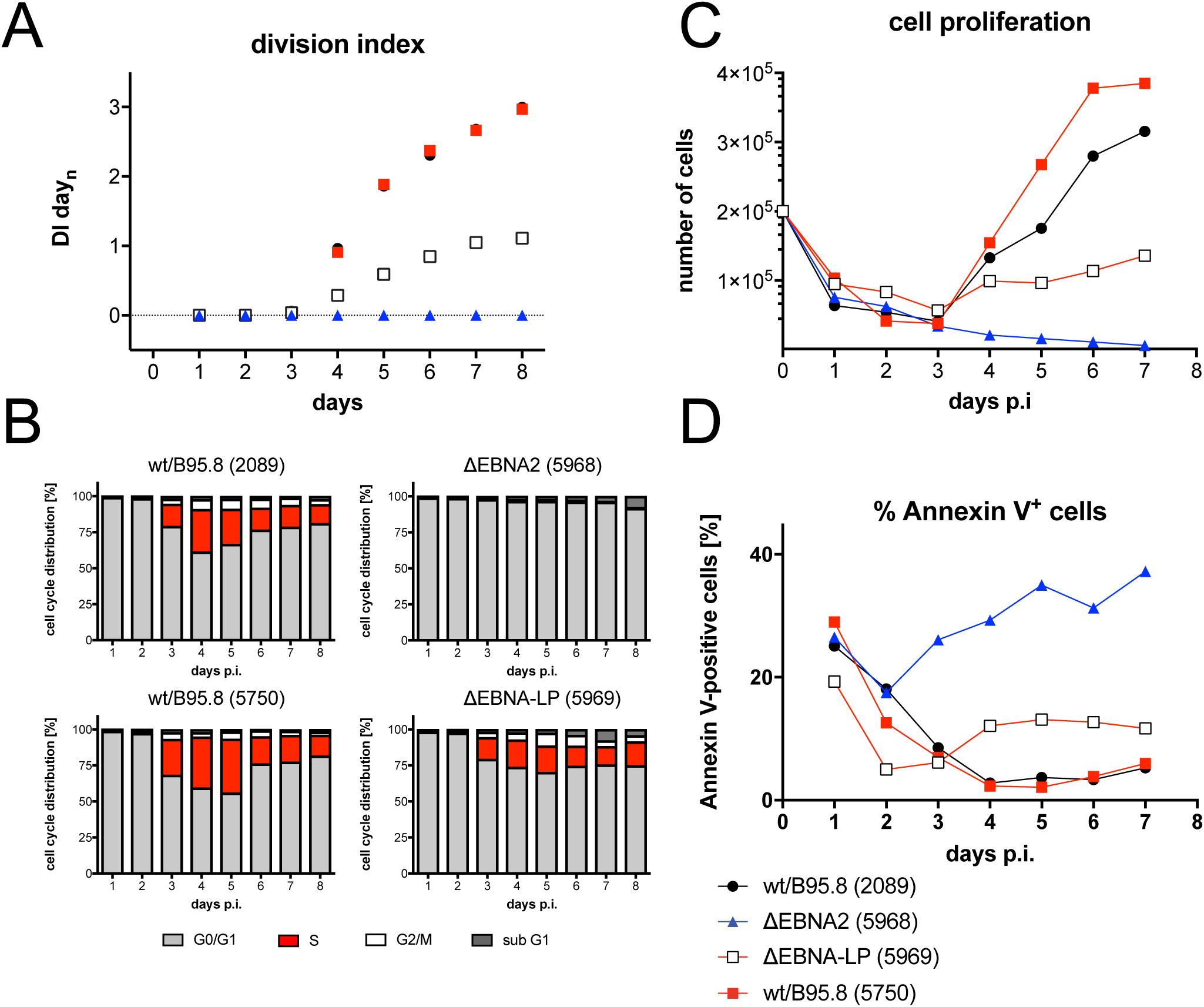
Activation kinetics of naïve B-lymphocytes infected with mutant EBVs negative for EBNA-LP or EBNA2. As in Fig. 3, sorted naïve B-lymphocytes were loaded with an intracellular dye and infected with the indicated viruses, which are wild-type with respect to viral latent genes (wt/B95.8 (2089) and wt/B95.8 (5750)) or are incapable of expressing EBNA2 (ΔEBNA2 (5968)) or EBNA-LP (ΔEBNA-LP (5969)). **(A)** Division Index is shown. **(B)** Cell cycle distributions of cultivated cells are provided. **(C)** Viable cells were counted by FACS. **(D)** Binding of Annexin V was analyzed by FACS. One representative experiment out of five experiments with B-lymphocytes from individual donors is shown.

### EBNA-LP has auxiliary, supportive functions in B cell activation and stable latent infection

The function of EBNA2 is well defined but the contribution of EBNA-LP to B cell activation and transformation is less clear. Early experiments with a C-terminally truncated EBNA-LP indicated that its functions might be critical in supporting the initial steps during EBV infection (Hammerschmidt and Sugden, 1989) and a recent report suggested that EBNA-LP is an essential gene for the transformation of naïve B-lymphocytes (Szymula et al., 2018). We generated a pair of recombinant EBVs with six complete copies of EBV’s BamHI-W-repeats each of which contains the W-exons of EBNA-LP (Fig. S2). The EBV recombinant wt/B95.8 (5750) is essentially wild-type (Tab.1), but its derivative ΔEBNA-LP (5969) is incapable of expressing EBNA-LP because each BamHI-W-repeat carries a translational stop codon in the W1 exon of EBNA-LP (Fig. S2).

Sorted naïve B-lymphocytes were infected with both viruses and the infected cells were analyzed daily. Cells infected with ΔEBNA-LP (5969) divided less vigorously (Fig. 5A) and showed an increased fraction of Annexin V-positive cells (Fig. 5D). DNA synthesis was first detectable on day 3 p.i., but the percentage of S-phase cells remained lower thereafter (Fig. 5B) compared with the two wild-type EBV stocks wt/B95.8 (2089) and wt/B95.8 (5750). The initial loss of B-lymphocytes during the first three days of infection was comparable (Fig. 5C). The first cell division was clearly detectable on day 4 p.i., but the ΔEBNA-LP (5969) infected cells did not support the rapid phase of cell proliferation as seen with the wild-type controls (Fig. 5A and C). As a consequence, the number of B cell infected with ΔEBNA-LP (5969) increased very slowly over time.

We also asked if naïve B-lymphocytes infected with ΔEBNA-LP (5969) can establish stable lymphoblastoid cell lines. From different anonymous donors, we expanded naïve B cells infected with ΔEBNA-LP (5969) to considerable cell numbers. We analyzed the protein expression of the established lymphoblastoid cell lines seven to eight weeks p.i. and found that the cells did not express EBNA-LP protein as expected (Fig. S3). However, once established these cells maintained their proliferative phenotype indistinguishably from wild-type EBV infected lymphoblastoid cell lines.

### EBV’s microRNAs support B cell reprogramming

EBV encodes up to 44 viral miRNAs that were found to support the survival of human B-lymphocytes in the early phase of infection (Seto et al., 2010; Feederle et al., 2011). We reanalyzed the phenotypes of sorted naïve B-lymphocytes infected with two mutant EBVs that express no viral miRNAs. The results confirmed the previous findings and documented that B cells infected with ΔmiR (4027) or r_ΔmiR (6338) (Tab. 1) showed a higher ratio of Annexin V-positive cells and accumulated to lower cell numbers in the first eight days of infection (Fig. 6 and Fig. S4). The reduced cell numbers were also due to a lower division index and a reduced percentage of S-phase cells compared with their parental wild-type EBV stocks confirming that EBV’s miRNAs regulate early cell cycle functions (Seto et al., 2010). Remarkably, the phenotypes observed with naïve B cells infected with the two ΔmiR mutants or the ΔEBNA-LP (5969) mutant EBV (Fig. 5) are comparable.

**Fig. 6.**
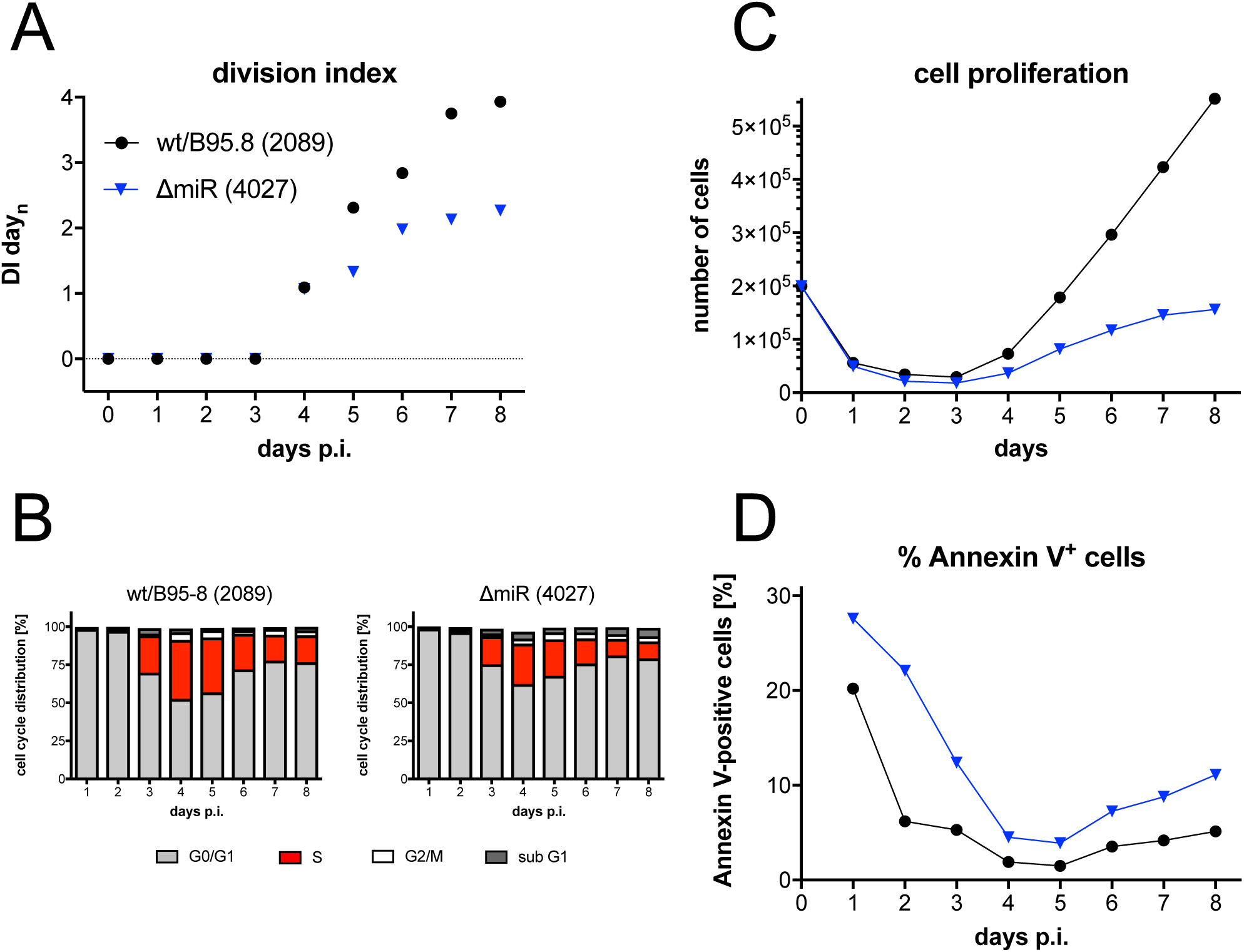
Activation kinetics of naïve B-lymphocytes infected with an EBV strain lacking all viral miRNAs. As in Fig. 3, sorted naïve B-lymphocytes were loaded with an intracellular dye and infected with wild-type wt/B95.8 (2089) EBV or ΔmiR (4027) negative for EBV’s miRNAs (Seto et al., 2010). Division Index is shown. **(B)** Cell cycle distributions of cultivated cells are provided. **(C)** Viable cells were counted by FACS. **(D)** Binding of Annexin V was analyzed by FACS. One representative experiment out of five experiments with B-lymphocytes from five individual donors is shown.

### LMP2A but not LMP1 nor the two non-coding EBERs contribute to the pre-latent phase

Apart from EBNA2, EBNA-LP and the miRNAs, other latent EBV genes that could play a critical role in the first days of infection include LMP1, LMP2A, EBER1, and EBER2. Their roles in activating and maintaining B cell proliferation have been studied by many groups, but their early functions are controversial. We infected sorted naïve B-lymphocytes isolated from several donors with the knock-out mutants ΔLMP1 (2597), ΔLMP2A (2525), and ΔEBER (6431) in which both EBER1 and EBER2 were deleted (Tab. 1) and found reproducible differences only in cells infected with ΔLMP2A (2525) compared with parental EBVs (Fig. 7, Fig. S4, Tab. 1). We also investigated a mutant EBV, ΔEBER/ΔmiR (6432), which is deficient in expressing the two viral non-coding RNAs EBER1 and 2 and all 44 miRNAs, because they might act in combination. Again, the phenotype of B cells infected with the double mutant did not differ from that of the ΔmiR (6338) mutant EBV (Fig. S4) documenting that only the viral miRNAs but not the two longer EBER RNAs play a discernable role in the pre-latent phase.

**Fig. 7.**
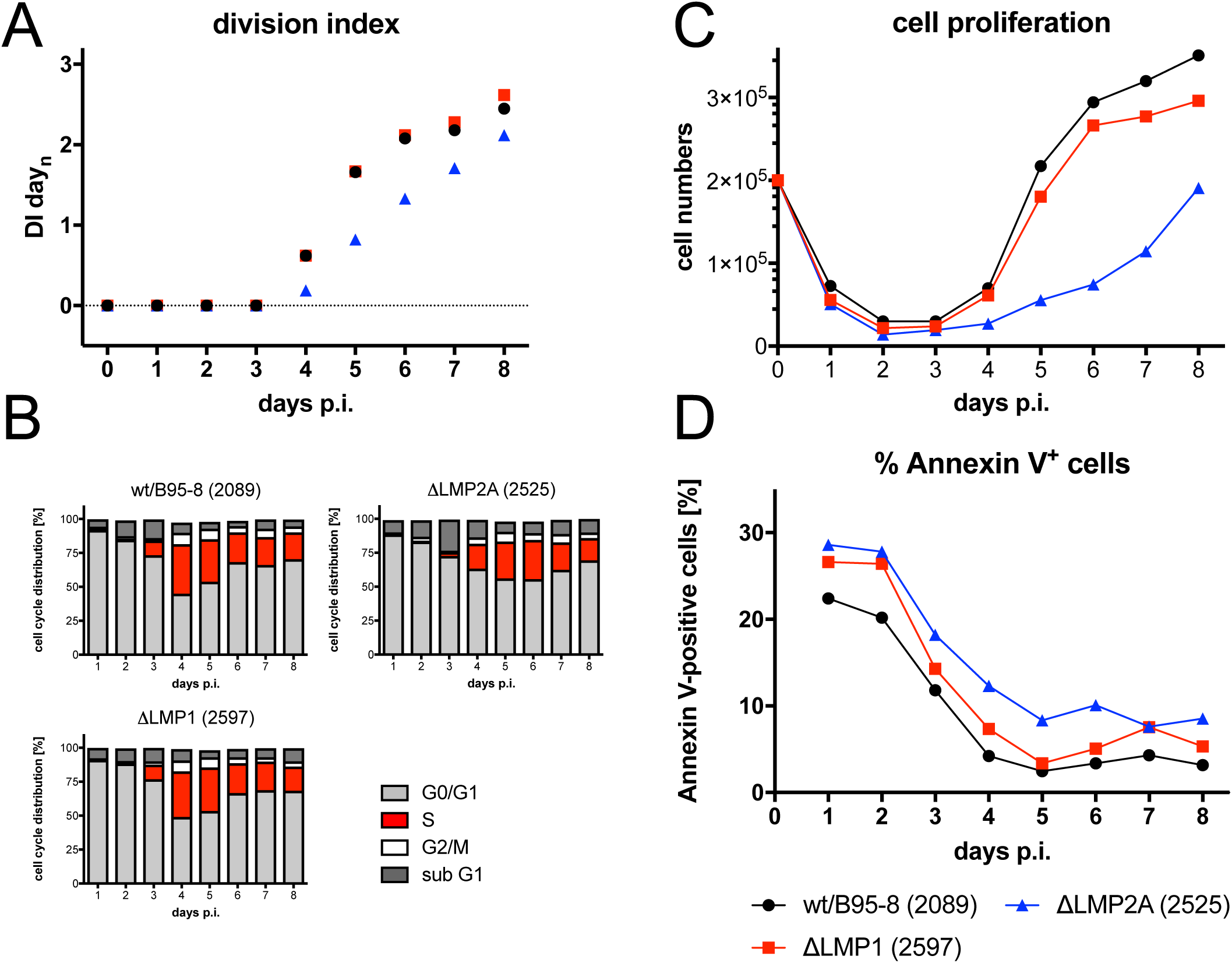
Activation kinetics of naïve B-lymphocytes infected with an LMP1-negative EBV strain. As in Fig. 3, sorted naïve B-lymphocytes were loaded with an intracellular dye and infected with the wild-type wt/B95.8 (2089) EBV strain or with ΔLMP1 (2597) EBV, which carries a knockout of EBV’s latent membrane protein 1, LMP1 (Dirmeier et al., 2003). **(A)** Division Index is shown. **(A)** Cell cycle distributions of cultivated cells are provided. **(C)** Viable cells were counted by FACS. **(D)** Binding of Annexin V was analyzed by FACS. One representative experiment out of three experiments with B-lymphocytes from three individual donors is shown.

### EBV mutants in EBNA3A and C show a proliferative advantage in the pre-latent phase

The early functions of EBNA3A and EBNA3C have been reported (Nikitin et al., 2010), but we wished to include these two latent genes in our study to explore the phenotypes of three single and double EBNA3A and 3C mutant EBVs (Tab. 1). With knock-out EBVs devoid of EBNA3A (6077) or EBNA3C (6123) we found no substantial differences compared with the parental wild-type EBV (Fig. 8), but noticed slightly higher cell numbers 5 to 6 days p.i.. This finding led us to engineer and test an EBV mutant deficient in expressing both viral genes. In several experiments with sorted naïve B-lymphocytes from different donors, the ΔEBNA3A/C (6331) mutant induced a slightly more robust proliferation and fewer Annexin V-positive cells compared with its parent (Fig. 8).

**Fig. 8.**
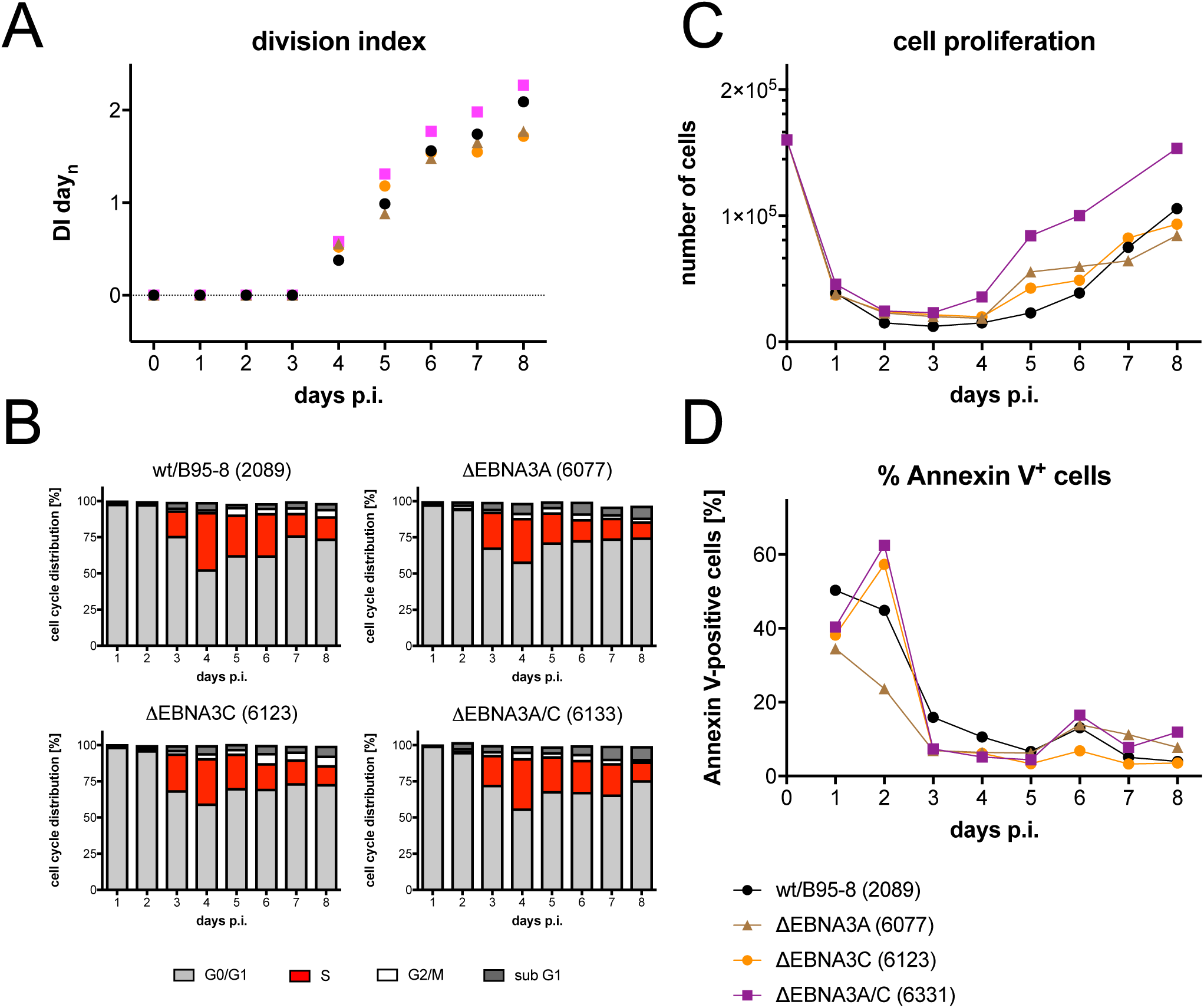
Activation kinetics of naïve B-lymphocytes infected with an EBV strains incapable of expressing EBNA3A, EBNA3C, or both EBNA3 family members. As in Fig. 3, sorted naïve B-lymphocytes were loaded with an intracellular dye and infected with a wild-type strain (wt/B95.8 (2089)) or three EBV strains deficient in EBNA3A, EBNA3B or both as indicated (Tab. 1). **(A)** Division Index is shown. **(B)** Cell cycle distributions of cultivated cells are provided. **(C)** Viable cells were counted by FACS. **(D)** Binding of Annexin V was analyzed by FACS. One representative experiment out of three experiments with B-lymphocytes from three individual donors is shown.

EBNA3C was reported to prevent a substantial DNA damage response (DDR) observed in the early phase of infection during cellular hyperproliferation in EBV-infected human B-lymphocytes (Nikitin et al., 2010). Given our findings so far, we wondered if the apoptotic loss of cells during the first three days of infection or later phenotypes might stem from this type of antiviral stress response that EBV infected B cells presumably experience prior to establishing viral latency. In naïve B-lymphocytes infected with wt/B95.8 (2089) EBV, we analyzed the phosphorylation of the histone variant H2A.X at Ser 139 (γ-H2A.X) by FACS over time. In addition to its well established role in the DDR for recognition and repair of double strand breaks, phosphorylation of H2A.X can also be triggered by DNA replication stress (DRS) during the surveillance of ongoing DNA replication in the absence of physical DNA damage (Ward and Chen, 2001; Saldivar et al., 2017).

Levels of γ-H2A.X were very low in sorted naïve and uninfected B-lymphocytes and in intact cells infected for the first two days (Fig. 9A). On day 3 and 4 p.i., the global levels of γ-H2A.X increased considerably in the population of viable cells and decreased again on day 5 p.i. and later (Fig. 9A). Interestingly, short term incubation with etoposide, a potent inducer of DDR, led to a considerable increase of γ-H2A.X staining in uninfected and infected cells indicating that the potential of EBV-infected cells to respond to DNA damage is intact in the pre-latent phase. We also analyzed B cells infected with ΔEBNA3A/C (6331). The cells showed marginally higher levels γ-H2A.X staining compared with wild-type EBV infected cells starting on day 6 p.i. (Fig. 9A), but clearly no major shift as seen after short term etoposide induction. Increased phosphorylation levels of H2A.X paralleled the onset of DNA synthesis on day 3 p.i. (Fig. 8B) and remained equally high during the initial, very rapid cell divisions on day 4, but were lower on the following days suggesting that γ-H2A.X staining reflects DRS rather than the cells’ response to DNA damage (Zeman and Cimprich, 2014).

**Fig. 9.**
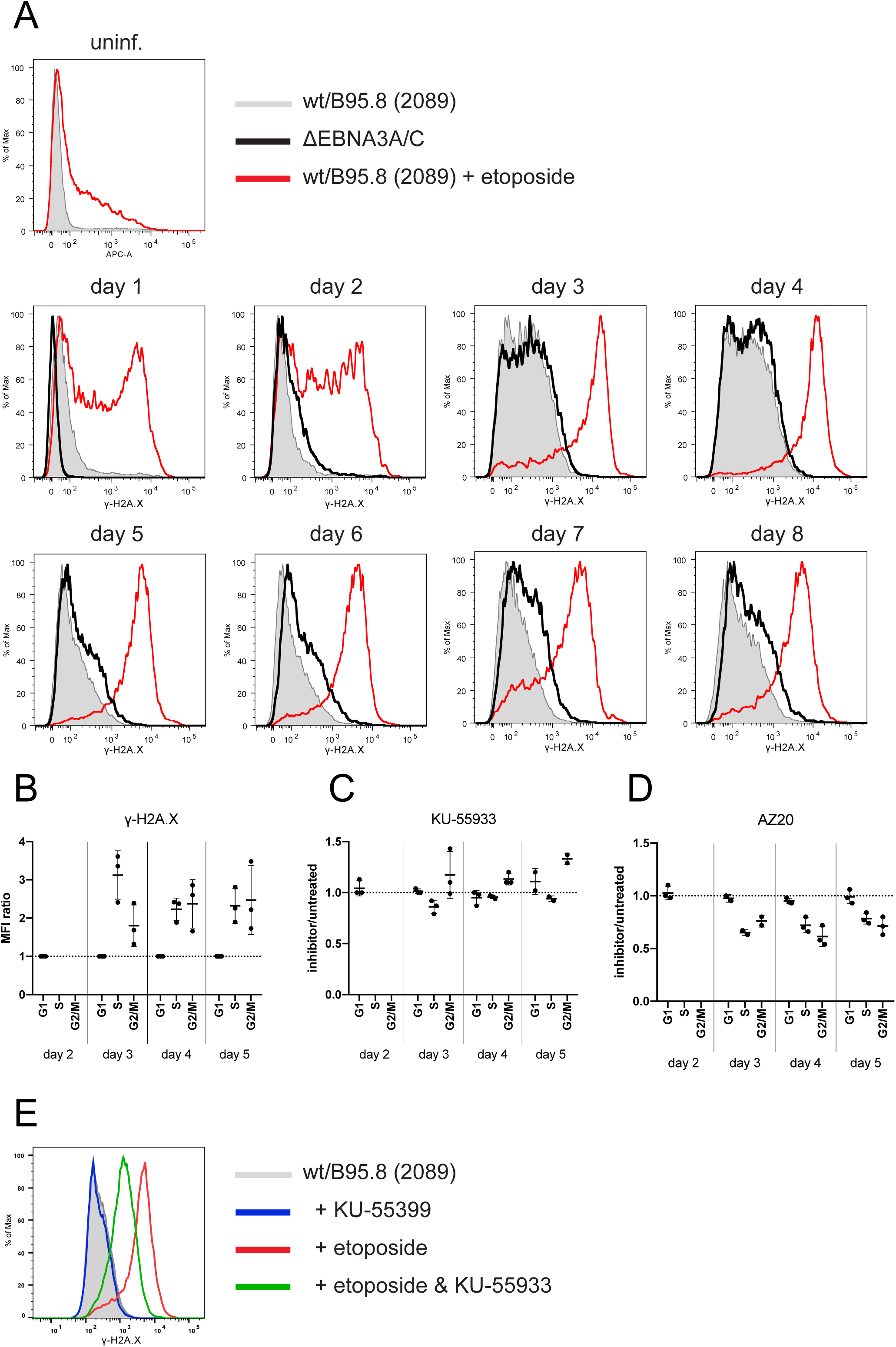
_γ_-H2A.X levels in uninfected B-lymphocytes and cells infected with wild-type EBV or an EBNA3A/3C double knockout EBV with ATM and ATR inhibitors. **(A)** FACS-sorted naïve B-lymphocytes were infected with wt/B95.8 (2089) EBV or ΔEBNA3A/C (6331) EBV with an MOIs of 0.1 and analyzed at different time points as indicated. Intracellular FACS staining detected the levels of the phosphorylated histone variant H2A.X (γ-H2A.X) in viable cells (according to scatter criteria by FACS). As a positive control, uninfected B-lymphocytes or cells infected with wt/B95.8 (2089) EBV were treated with 85 μM etoposide for 1 h prior to harvest (red line). One representative experiment out of three with B-lymphocytes from three individual donors is shown. **(B)** FACS-sorted naïve B-lymphocytes were infected with B95.8 EBV with an MOIs of 0.1. The cells were pulse-labeled with EdU (5-ethynyl-2’-deoxyuridine, a nucleoside analog to thymidine incorporated into DNA during active DNA synthesis) for one hour, fixed, permeabilized, and frozen at −80°C at the indicated time points. For FACS analysis, the cells were thawed and analyzed after intracellular staining with an antibody directed against γ-H2A.X and a click reaction between EdU and Alexa Fluor 488. Single viable cells were considered by gating and the mean fluorescence intensities (MFI) of the fluorochrome coupled γ-H2A.X specific antibody were separately recorded in the cell cycle fractions G1, S, and G2/M. MFI values of the S and G2/M fractions were compared with the MFI values of G1 cells and expressed as ratios of S versus G1 or G2/M versus G1 MFI levels. MFI levels of γ-H2A.X fluorescence in G1 were set to 1.0. On day 3 to 5, cells in S and G2/M had two to three-fold higher γ-H2A.X levels compared with cells in G1. Cells infected for two days do not cycle or synthesize DNA. Three independent experiments are summarized. **(C)** B95.8 infected B-lymphocytes were incubated with EdU together with the ATM inhibitor KU-55933, or were incubated with EdU, only, for one hour prior to harvest as in panel B. Shown are the MFI ratios of γ-H2A.X levels in the different cell cycle fractions of KU-55933 treated versus untreated cells. KU-55933 showed a slight inhibitory effect on day 3 p.i. in S phase cells, but other γ-H2A.X levels were not reduced but sometimes even elevated when KU-55933 was applied. Three independent experiments are summarized. **(D)** The experimental setup is identical to panel C except that the ATR inhibitor AZ20 was applied for one hour together with EdU. Cells in S and G2/M but not cells in G1 showed a clear reduction of γ-H2A.X levels in the presence of AZ20 on day 3, 4, and 5 p.i.. Three independent experiments are summarized. **(E)** The histogram demonstrates the inhibitory effect of the inhibitor KU-55933 on an etoposide-induced DDR. B-lymphocytes were infected with wt/B95.8 (2089) EBV for five days and treated with KU-55933 for one hour prior to analysis with the γ-H2A.X specific antibody by intracellular FACS (red line in the histogram). Concomitant addition of the ATM inhibitor KU-55933 for 1 hour together with etoposide reduced the induced levels of γ-H2A.X considerably (green), whereas the addition of KU-55933 alone (blue) had no effect on cells that were not treated with etoposide (grey shaded histogram).

Both genotoxic and non-genotoxic stresses interrupt the p53-MDM2 loop to stabilize p53 increasing its steady state protein level. This process leads to changes in the expression of hundreds of p53-responsive genes including the check point inhibitor *WAF1/CIP1* coding for p21, and induces a halt in proliferation preventing the transmission of damaged DNA to daughter cells. We looked at the protein levels of both p53 and p21 as well as Ku70 and Rad51, which are commonly upregulated during DDR. We found considerable levels of p53 and p21 in wild-type EBV infected cells starting on day 4 p.i. and onwards (Fig. S5) and, in addition, levels of Ku70 and Rad51 also increased in parallel with p53 and p21. Unexpectedly, in B cells infected with ΔEBNA3A/C (6331) we barely detected p53 protein and found slightly reduced levels of p21 compared with wt/B95.8 (2089) infected cells (Fig. S5). Levels of Rad51 appeared marginally reduced in cells infected with ΔEBNA3A/C (6331) whereas Ku70 levels seemed to be similar in cells infected ΔEBNA3A/C (6331) or wt/B95.8 (2089) EBV. The findings suggest that cells infected with the EBNA3A/EBNA3C double mutant EBV might experience lower stress levels, which could explain their proliferating more vigorously than wild-type EBV-infected B-lymphocytes in the pre-latent phase (Fig. 8).

### In the pre-latent phase EBV infection does not induce a DNA damage response but causes DNA replication stress

We were puzzled by these unexpected findings because they are in obvious conflict with a previous report (Nikitin et al., 2010). To discriminate between DDR and DRS as the confounding source of the elevated γ-H2A.X signal, we asked in which phase of the cell cycle the infected cells show an increase in γ-H2A.X levels. We reasoned that if DRS is the major driver of this effect, it should only increase γ-H2A.X levels in S-phase and post-replicative G2/M cells, whereas DDR would equally affect all cells independent of the cell cycle state. We infected sorted naïve B-lymphocytes from adenoid tissue with the reference strain B95.8 and an MOI of 0.1 and labeled the newly synthesized cellular DNA with EdU (5-ethynyl-2’-deoxyuridine), a nucleoside analog to thymidine for one hour prior to analysis. We found that cells in S and in G2/M showed a two to three-fold higher level of γ-H2A.X staining, whereas G1 phase cells were unaffected (Fig. 9B). To further support our hypothesis, we combined this analysis with two inhibitors, KU-55933 (Hickson et al., 2004) or AZ20 (Foote et al., 2013), known to specifically block activation of ATM (ataxia telangiectasia-mutated) and ATR (ATM and rad3-related), the two major DNA damage checkpoint kinases responsible for γ-H2A.X phosphorylation. Whereas ATM is primarily activated by DDR, ATR responds to stretches of RPA-coated ssDNA at stalled replication forks during DRS (Cimprich and Cortez, 2008). While KU-55933 clearly repressed an etoposide induced DDR signal in EBV infected B cells (Fig. 9E), it did not reduce the mean fluorescence intensity (MFI) of γ-H2A.X staining in EBV infected cells on day 3 and 4 p.i. but merely increased it in S and G2/M phase cells (Fig. 9C). In contrast AZ20 caused a considerable reduction of γ-H2A.X levels in all actively cycling cells on day 3 to 5 (Fig. 9D), indicative of a reduction in the DRS as the primary source of the elevated γ-H2A.X signal.

### EBNA1 does not contribute to B cell activation, cell cycle entry or early cell proliferation

Next to EBNA2, EBNA1 is the second latent protein, whose functions have been studied intensively. EBNA1 is critical for the extrachromosomal maintenance of genomic EBV DNA and it acts as a transcription factor regulating the expression of viral and cellular genes in latently EBV-infected cells. We asked whether this viral factor makes important contributions during early infection and engineered a viral mutant, in which the translation of EBNA1’s open reading frame is disabled. Sorted naïve B-lymphocytes were infected with this mutant EBV, termed ΔEBNA1 (6285), or with the reference wild-type strain wt/B95.8 (2089). Surprisingly, the phenotypes of B cells infected with this pair of viruses differed barely. Cells infected with ΔEBNA1 (6285) or wt/B95.8 (2089) underwent similar rates of initial death, became similarly activated, started cellular DNA synthesis on day 3, and began with rapid cell divisions on day 4 p.i. as we had observed with all mutant EBVs but ΔEBNA2 (5968) so far (Fig. 10). Proliferation of B cells infected with ΔEBNA1 (6285) slowed down starting on day 6 or 7 p.i. (depending on the donor’s B cells) but showed only a slight increase in Annexin V-positive cells (Fig. 10B). We analyzed the protein levels of EBNA2, EBNA1, and MYC and found that cells infected with ΔEBNA1 (6285) expressed no EBNA1, as expected, but comparable levels of EBNA2 and MYC protein on day 4. p.i. (Fig. 10E). On day 8 p.i., however, the levels of both EBNA2 and MYC were considerably lower than wild-type EBV-infected cells, but B cells infected with ΔEBNA1 (6285) survived even longer, shrank in volume (Fig. S6) and became extinct about two to three weeks post infection. Whereas EBNA1 plays a central role in the maintenance of EBV latency, or data indicate that EBNA1 is absolutely dispensable for the early stage of EBV-infected primary naïve B cells.

**Fig. 10.**
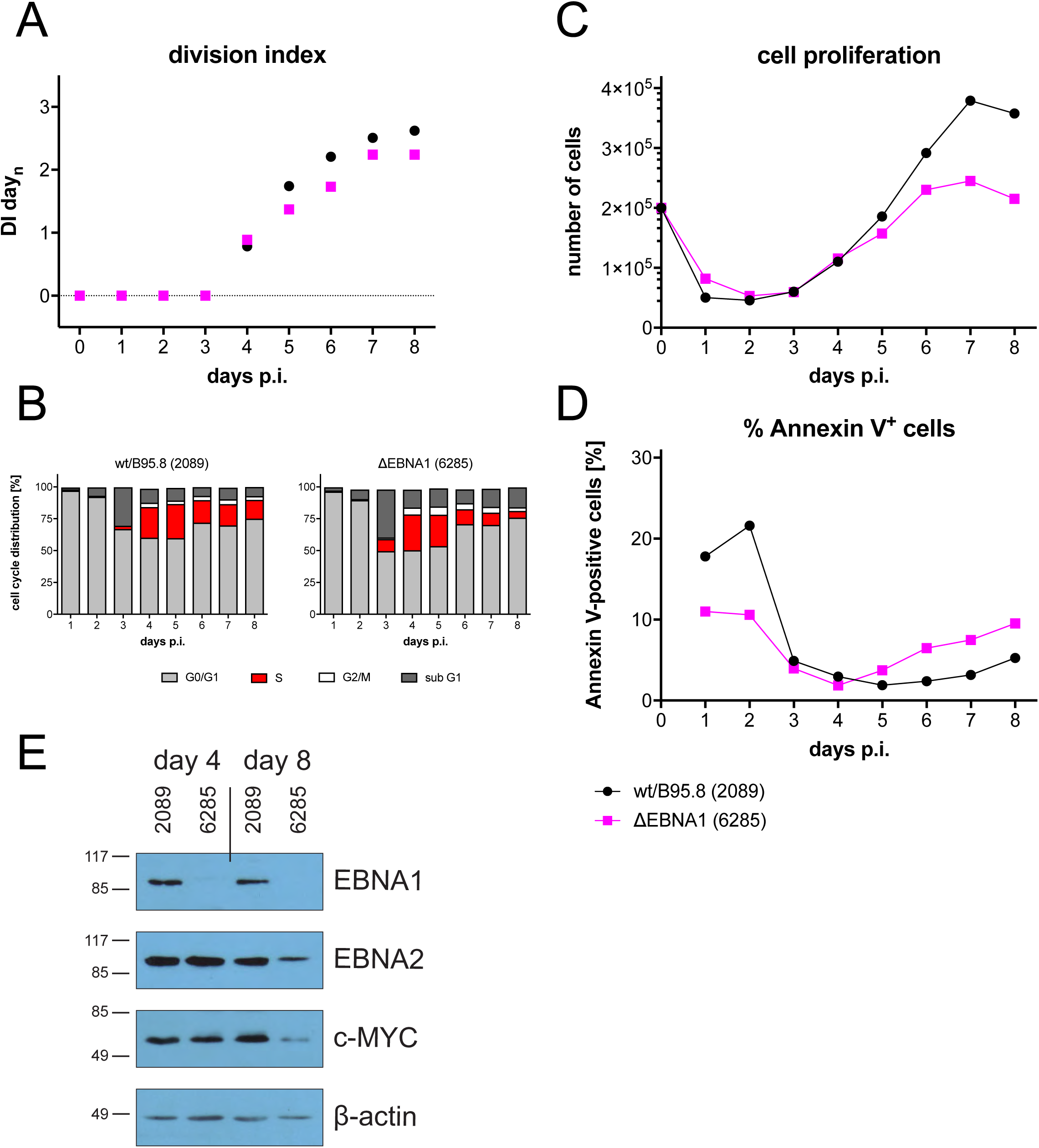
Activation kinetics of naïve B-lymphocytes infected with an EBV strain lacking EBNA1. As in Fig. 3, sorted naïve B-lymphocytes were loaded with an intracellular dye and infected with wild-type wt/B95.8 (2089) EBV or ΔEBNA1 (6285), which cannot express EBNA1 due to a point mutation in EBNA1’s translational start codon. **(A)** Division Index is shown. **(B)** Cell cycle distributions of cultivated cells are provided. **(C)** Viable cells were counted by FACS. **(D)** Binding of Annexin V was analyzed by FACS. **(E)** Steady state protein levels of B cells infected with wtB95.8 (2089) or ΔEBNA1 (6285) for 4 or 8 days were analyzed by Western blot immunodetection with antibodies directed against EBNA1, EBNA2, c-MYC, or ß-actin. One representative experiment out of five is shown.

### A minimal set of EBV genes is sufficient to support B cell activation and proliferation

So far, we learnt that many viral latent genes are dispensable in EBV infected B cells during the pre-latent phase. We wondered if the expression of only EBNA2, EBNA-LP, and BHRF1 might be sufficient to activate primary B-lymphocytes and drive them into proliferation. Towards this end we revisited the two EBV plasmids p554 and p613 we had engineered earlier (Hammerschmidt and Sugden, 1989). They both encompass a contiguous fragment of EBV with the nucleotide coordinates #7315 to #56,083 of the wt/B95.8 strain. This fragment carries *oriP*, EBNA-LP and EBNA2 but only two complete and a truncated copy of the BamHI-W-repeat element. This EBV DNA fragment also contains the lytic origin of DNA replication, *oriLyt*, together with the BHRF1 and BHLF1 loci, which flank it. The plasmid p554 and p613 DNAs can be packaged into infectious EBV particles, because p554 and p613 also contain TRs, the essential DNA packaging elements of EBV (Hammerschmidt and Sugden, 1989; Zimmermann and Hammerschmidt, 1995). p554 and p613 differ in that p613 is incapable of expressing EBNA2 (Hammerschmidt and Sugden, 1989).

We separately introduced the two plasmids into EBV particles with the help of a packaging cell line that harbors a non-transforming EBV helper virus genome with several genetic modifications that include a deleted EBNA2 locus (Hettich et al., 2006). Concentrated vector stocks with p554 or p613 were used to infect naïve B-lymphocytes and living cells (according to FACS forward and sideward scatter criteria) were analyzed for their Annexin V binding, TMRE staining (a marker of active mitochondria), and BrdU incorporation (Fig. 11). The vector stocks did not nearly reach EBV titers that were used in all the previous experiments (up to 4×10^6^ GRU/ml), but B cells infected with p554 clearly were alive on day 6 p.i. (Fig. 11). Only a small percentage of the living cells bound Annexin V (Fig. 11A), but a high fraction stained with TMRE indicating robust mitochondrial activity (Fig. 11B). These observations were in contrast to cells incubated with p613 vector stocks devoid of EBNA2 (Fig. 11B). At a much lower level than wt/B95.8 (2089) EBV infected cells B cells infected with p554 vector stocks incorporated BrdU indicating that they underwent DNA replication, mitosis, and cell divisions (Fig. 11C). On day 6 p.i., few intact B cells infected with p613 vector stocks or non-infected cells survived, which did not cycle (Fig. 11C). This experiment confirms that EBNA2 is the essential viral factor in the pre-latent phase of human B cells and suggests that it probably in conjunction with EBNA-LP might be sufficient to reprogram resting human B-lymphocytes.

**Fig. 11.**
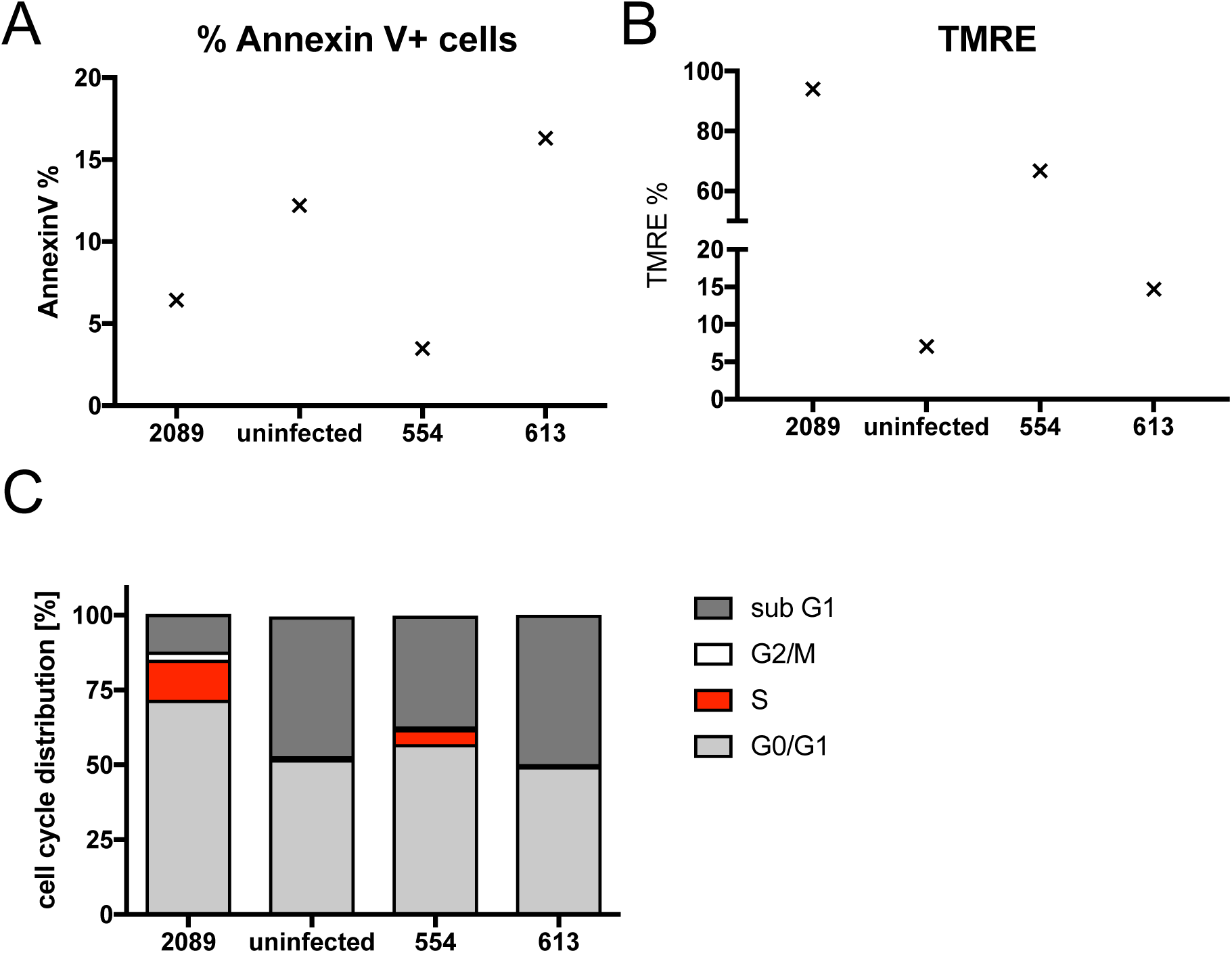
A vector approach identifies EBNA2 as an essential viral gene in EBV’s pre-latent phase. Naïve primary B-lymphocytes were infected for six days with vector stocks obtained by packaging the plasmids p554 and p613 (Hammerschmidt and Sugden, 1989) into EBV-based viral particles (Hettich et al., 2006). Non-infected B-lymphocytes and cells infected with an MOI of 0.1 of wt/B95.8 (2089) virus stocks served as negative and positive controls, respectively. Annexin V binding and TMRE staining of active mitochondria indicated non-apoptotic and metabolically active cells, respectively. BrdU incorporation revealed cycling cells in S phase. One representative experiment out of three is shown. p554 encodes EBNA2, EBNA-LP, BHRF1 and BHLF1, whereas p613 lacks EBNA2.

## Discussion

### Reconstruction of a B95.8 based EBV field strain

In this study, we present a version of the B95.8 strain of EBV (Miller and Lipman, 1973) in which we reverted the ∼12 kb deletion present in this laboratory isolate (Bornkamm et al., 1980). Recently, another group has followed this approach (Kanda et al., 2015) using a fragment from the Akata strain of EBV. The authors demonstrated that the miRNAs absent in B95.8 were expressed in cells infected with the reconstituted virus. They also showed a downregulation of the *NDRG1* gene expression in epithelial cells infected with the reconstituted virus as compared to cells infected with their recombinant parental B95.8 strain (Kanda et al., 2015). We employed our BAC-derived wt/B95.8 (2089) strain to obtain a virus that is as close as possible to a B95.8 based field strain and to allow further engineering and reverse genetic experiments. B-lymphocytes infected with the reconstituted strain r_wt/B95.8 (6008) or the parental wt/B95.8 (2089) strain behave very similarly in our proliferation experiments (Fig. 3), but we suspect that the presence of all BART miRNAs in r_wt/B95.8 (6008) will lead to observable phenotypes in other infection models. As we have shown recently the viral miRNAs play a remarkable role in inhibiting the T cell recognition of EBV infected cells *in vivo* and *in vitro* (Murer et al., 2019; Albanese et al., 2016; Tagawa et al., 2016). It will be interesting to compare both related viruses in e.g. humanized mice that may reveal important additional roles of the BART miRNAs encoded in the r_wt/B95.8 (6008) strain of EBV, only.

### The limitations of the in vitro B cell infection model

When we started our experimental work, we did not expect to learn that EBNA2 is the only viral latent gene product that is essential (and presumably also sufficient) to activate resting human primary B cells and to induce their cycling. EBNA-LP, LMP2A, and the viral miRNAs contribute to this process, but naïve B-lymphocytes infected with mutant EBVs that do not express these individual auxiliary genes readily give rise to lymphoblastoid cell lines. In this experimental model even EBNA1 seems to be dispensable for several days but cells infected with an EBNA1-negative EBV only very rarely yield stable cell lines in which the EBV genome is then found chromosomally integrated (Humme et al., 2003). We also learnt that the newly infected cells express very high levels of EBNA2 initially (Fig. 12A, Mrozek-Gorska et al., 2019), which now appear sufficient to reprogram the infected cells until other latent viral gene products support and stabilize the reprogrammed cells long-term. Such a secondary role has been proposed for LMP1 (Price et al., 2012). LMP1’s mRNA is expressed at substantial levels early on (Mrozek-Gorska et al., 2019), but an LMP1-negative EBV mutant did not result in a distinct phenotype in the pre-latent phase (Fig. 7), confirming recent reports (Price et al., 2012; Price et al., 2018).

**Fig. 12.**
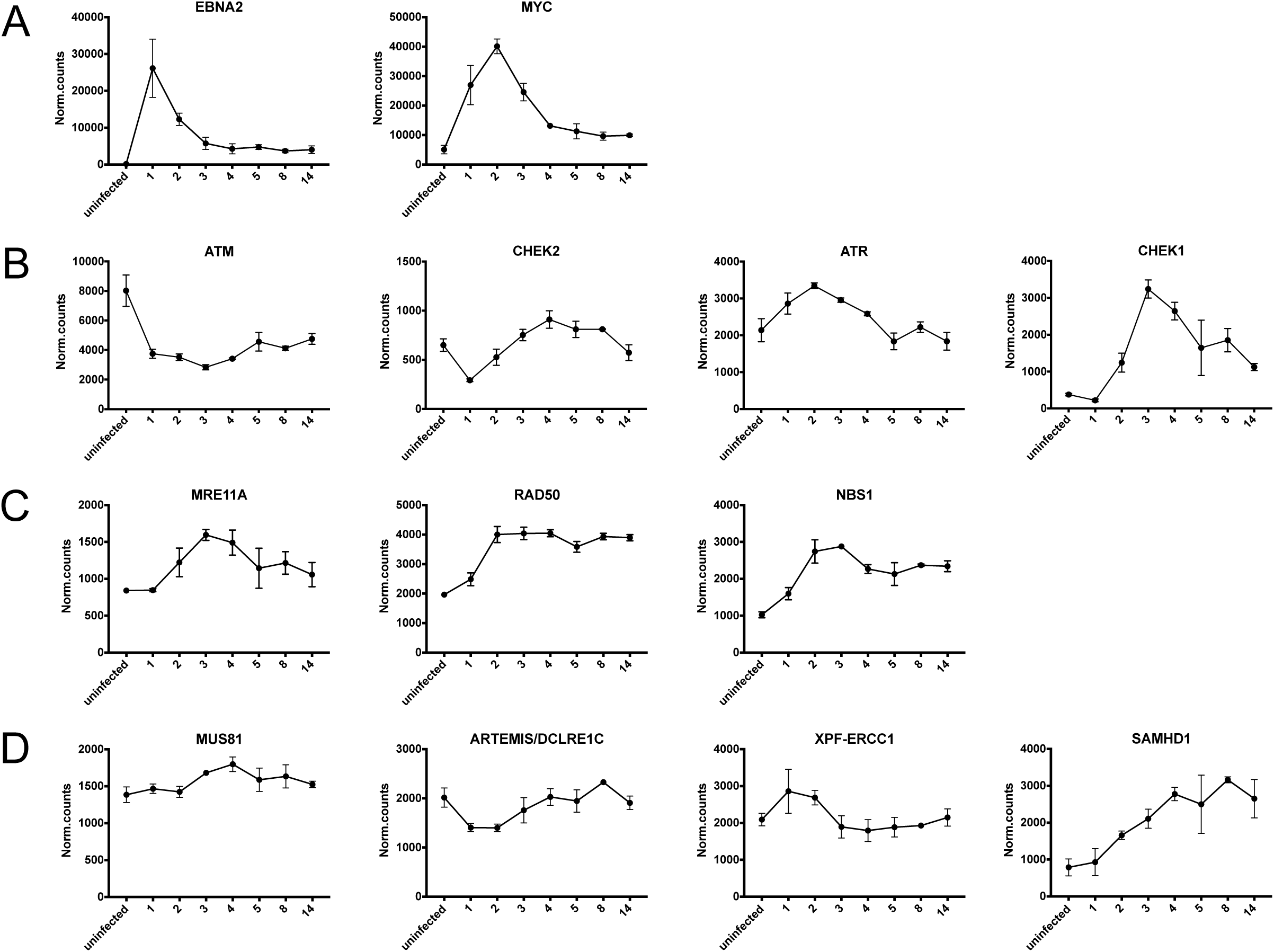
Transcriptional regulation of EBNA2 and selected cellular genes involved in DNA damage response or DNA replication stress. Primary naïve B-lymphocytes were isolated and infected with wt/B95.8 (2089) EBV. The transcriptomes of uninfected and infected B cells were investigated by RNA-seq technologies at the indicated time points as described previously (Mrozek-Gorska et al., 2019). **(A)** EBNA2 and MYC appear to be coregulated. **(B)** The transcriptional regulation of the ATM-Chek1(Chk1) and ATR-Chek2(Chk2) pathways, which are activated by DNA double-strand breaks and single-stranded DNA, respectively, is shown. **(C)** The transcriptional regulation of components of the MRN complex and **(D)** genes involved in double-strand break and single-strand DNA repair, respectively, is shown. The data were obtained from our recent work (Mrozek-Gorska et al., 2019) and are freely accessible (https://scialdonelab.shinyapps.io/EBV_B/).

We found that the ratio of EBV versus its target cell is critical and can even appear toxic at high doses (Fig. 1). At our standard dose (MOI of 0.1) we can safely assume that each B-lymphocyte is infected with at least one biologically active EBV particle on average given our previous investigations with Daudi cells (Steinbrück et al., 2015). A higher fraction of B-lymphocytes can be identified initially to be EBNA2-positive, when the virus dose is increased (Fig. 1C), but the effect is counterproductive and reduces the number of latently infected cells 8 days p.i. (Fig. 1A). An MOI of 3.0, which equals at least 30 biologically active virions per B-lymphocytes, is acutely toxic to the cells (Fig. 1B), suggesting that many copies of genomic EBV DNA lead to a high gene dose of EBNA2, which seems to be detrimental. Very high levels of EBNA2 protein likely result in extreme MYC levels that induce ARF expression (Ko et al., 2018), which in turn activates p53 and leads to cellular senescence, cell cycle arrest, and apoptotic death (Zindy et al., 1998). This working hypothesis seems plausible but needs to be tested.

The prominent function of EBNA2 in this *in vitro* infection model has been known for a long time, but EBNA2 is rarely found expressed in cells *in vivo* (Longnecker et al., 2013). Even EBV field strains in which the EBNA2 gene is deleted have been identified in latently infected Burkitt lymphoma cells, a rare finding (Kelly et al., 2002; Kelly et al., 2006). How these EBV strains evolve is speculative, but it is likely that the deletion of EBNA2 is a late event in these cells, which are under control of the *MYC* oncogene translocated into an active immunoglobulin locus.

In mice reconstituted with functional components of a human immune system it is possible to study EBV’s biology *in vivo* (Münz, 2015), but steps early after infection are difficult to investigate in this tractable model. To our knowledge, an EBNA2-deleted virus has not been studied in this animal model, yet, but, unexpectedly, EBV strains incapable of expressing EBNA3C were found to establish latency and cause B cell lymphomas albeit at a reduced frequency compared with wild-type EBV (Murer et al., 2018; Romero-Masters et al., 2018). EBNA3C is a transcriptional repressor and has been found to abrogate the expression of cell cycle inhibitors such as p16^INK4A^ encoded by *CDKN2A*, which is a prerequisite to establish lymphoblastoid cell lines with EBNA3C-deleted EBVs *in vitro* (Skalska et al., 2013). Our ΔEBNA3C (6123) mutant EBV did not reveal an early phenotype compared with wild-type EBV infected B-lymphocytes and the cells showed robust activation and proliferation in the pre-latent phase (Fig. 8). B cells infected with EBNA3C deleted mutant EBVs ceased to proliferate about two weeks post infection (Skalska et al., 2013) very much in contrast to the situation *in vivo*. In hu-mice EBNA3C deleted viruses established latent infection in B cells while the cells expressed appreciable levels of p16 (Romero-Masters et al., 2018). Possibly, there are several reasons that may lead to the observed differences, but the comparison demonstrates the limitations of the experimental models with which we can study the biology of EBV infection *in vitro*.

Some of these limitations also became obvious in our experiments. Naïve primary B-lymphocytes isolated from many different donors showed certain unstable phenotypes when infected with EBV. For example, in the experiments shown in Figs. 1, 3 and 5, the numbers of B cells from two donors infected with different wild-type EBV strains exceeded the initially infected and seeded cell numbers on day 8 p.i.. The examples shown in Figs. 8 and S1 illustrate that B cells from two other donors showed a less robust recovery with respect to cell numbers when infected with wt/B95.8 (2089) EBV. In almost all settings at least three independent experiments with B cells from different donors were conducted to validate the experimental outcome with the mutant EBVs, but individual experiments showed a certain variability in particular with respect to cell numbers in the time course experiments. We have no firm clue about the molecular basis of this observation but believe that the experiments reflect both technical as well as donor specific conditions.

### The master regulator EBNA2

Several arguments underscore EBNA2’s prominent role in the pre-latent phase of infected primary B cells. EBNA2 is a viral enhancer factor and a potent inducer of the *MYC* gene (Kempkes et al., 1995c; Zhao et al., 2011). MYC is the master regulator of all cells and controls cellular metabolism, cell growth and differentiation, and cell cycle progression, but also governs apoptosis and cell death. It is a potent oncoprotein that is highly expressed in Burkitt lymphomas. High levels of MYC protein induce cellular growth, cell cycle entry and cell proliferation, but, as a consequence, also cause replication stress (Rohban and Campaner, 2015). Induced overexpression of *MYC* increases the fraction of S-phase cells and accelerates S-phase progression probably by initiating premature origin firing. This leads to a higher density of active origins of DNA replication, which are associated with asymmetrical fork progression and stalled replicative DNA intermediates (Srinivasan et al., 2013). The very high initial expression of EBNA2 already on day one p.i. (Fig. 12A, Mrozek-Gorska et al., 2019) probably causes the timely expression of *MYC* to reach its highest level on day 2 p.i. (Fig. 12A), which in turn explains most of the phenotypic changes we observed during the first days of EBV infected B cells.

EBNA2 directly targets *MYC* and activates its expression (Kaiser et al., 1999). EBNA2 is known to bind to two B cell enhancers about 500 kbp upstream of the *MYC* gene and brings them in contact with the *MYC* promoter element to achieve its transcriptional activation (Zhao et al., 2011). EBNA2 was also identified to be the key component of this enhancer complex together with additional cellular and viral factors such as NF-kB subunits and EBNA-LP, respectively (Zhou et al., 2015). To our knowledge, there are no data that describe the action of EBNA2 at these cis-acting elements during the very first days of infection, but our data suggest that EBNA2 acts on the *MYC* locus and its upstream enhancers initially, when EBV infects quiescent primary B-lymphocytes.

### Does infection with EBV induce a genotoxic signal in primary B cells?

When cellular machines sense damaged DNA, they induce a DNA damage response (DDR) and initiate DNA repair to prevent genetic instability of the cell. Similarly, during cellular DNA replication, replication forks can stall, when they encounter sequences that are intrinsically difficult to replicate or when the cell experience genotoxic stress during mitosis. These apparent problems of cells progressing through the S-phase of cell cycle are generally referred to as DRS. Many cellular factors are known that manage DDR and/or DRS to maintain the genetic integrity of the cell or to induce its subsequent death. Using our recent data set (Mrozek-Gorska et al., 2019) we asked whether EBV infection regulates these factors in the pre-latent phase to evaluate if they play a decisive role in the reprogramming and survival of the virally infected B cells. It is also obvious from these data that EBV infection regulates the gene expression of the ATM-Chk2 and ATR-Chk1 pathways, which are induced by DNA double-stranded breaks and exposed single-stranded DNA and cope with DDR and DRS, respectively (Fig. 12B). In particular Chk1 is dramatically induced starting on day 2 p.i. (Fig. 12B). The ATR-Chk1 pathway is active during DNA synthesis and acts primarily as a replication checkpoint.

The trimeric MRN complex, which consists of MRE11A, RAD50, and NBS1 (also called Nibrin or NBN), is a multifunctional DDR machine (Syed and Tainer, 2018). Together with other damage recognition receptors the MRN complex can detect damaged DNAs to initiate DNA repair signals but it can also start a cascade of innate immune signals when certain viral DNAs are detected. Not surprisingly DNA viruses such as adenovirus (Shah and O’Shea, 2015) target the MRN complex to prevent its functions during lytic infection with this pathogen, which induces a strong DDR signal when the amplified viral DNA accumulates (Lilley et al., 2007). EBV does not start its lytic phase initially as adenoviruses do (Hammerschmidt, 2015), but EBV infection upregulates the transcript levels of all three components of the MRN complex two to three-fold as early as on day two (Fig. 12C) and prior to the onset of cellular DNA synthesis one day later. The levels of RAD50 transcripts remain unchanged thereafter whereas MRE11A and NBS1 are slightly reduced when established lymphoblastoid cells emerge (Fig. 12C). It thus appears that these three proteins are upregulated prior to or concomitant with the first occurrence of DNA replication, but their mRNA levels remain constant or are reduced later when the cells start to proliferate rapidly on day 4 and 5 p.i..

The endonucleases, MUS81, ARTEMIS (DCLRE1C), and XPF-ERCC1 act on stalled replication forks to resolve DRS during the S and G2 phases and/or during mitosis and cleave single-stranded DNAs (Bétous et al., 2018). In the pre-latent phase, the transcript levels of these important enzymes vary modestly with the exception of XPF-ERCC1, which peaks on day one and two p.i. (Fig. 12D). In contrast, EBV infection induces SAMHD1, which promotes the degradation of nascent strand DNA at stalled replication forks by binding to and stimulating the exonuclease activity of MRE11A which is upregulated to reach maximal levels on day 3 p.i. (Fig. 12C). MRE11A degrades nascent strand DNA during fork resection and thus facilitates fork restart. Transcript levels of SAMHD1 increase over time and stay elevated when the cells undergo rapid rounds of cell division and DNA replication (Fig. 12D) suggesting that high levels of both MRE11A and SAMHD1 limit DRS (Coquel et al., 2018) in this phase of viral infection and later when lymphoblastoid cell lines emerge.

These additional data suggested that EBV infection does not globally induce cellular genes involved in DNA damage signaling or repair early after infection as reported previously (Nikitin et al., 2010), but rather selectively regulates genes with dedicated functions in DRS such as SAMHD1, Chk1, and MRE11A. This interpretation is supported by our experimental findings with the histone variant H2A.X, which is highly phosphorylated at serine 139 on day 3 and 4 (Fig. 9A), when the cells undergo the first round of DNA synthesis and rapid cell divisions, respectively. Primarily, phosphorylation of H2A.X is a sensitive marker for DNA double-strand breaks, which is the canonical function of γ-H2A.X with a central role in DDR (Rogakou et al., 1998; Shroff et al., 2004). Yet, phosphorylation of H2A.X also serves other functions for example in embryonic stem cells, during cellular senescence, and in X chromosome inactivation (Turinetto and Giachino, 2015 for a recent review). In cycling cells phosphorylation of H2A.X increases as cells progress from G1 into S, G2 and M phase, reaching maximal levels at metaphase in the absence of DNA damage (McManus and Hendzel, 2005; Tu et al., 2013; Ward and Chen, 2001). H2A.X phosphorylation might have a role in regulating the activation of the mitotic spindle assembly checkpoint (SAC) or in the formation of an intact mitotic check point complex during mitosis (Turinetto and Giachino, 2015) documenting that cycling cells show elevated but physiological levels of γ-H2A.X. The DNA damage checkpoint kinase ATR is always active during each S-phase, which is due to longer stretches of single-stranded DNA that inherently activate ATR (Berti and Vindigni, 2016) to orchestrate the resolution of stalled replication forks to avoid DNA damage, eventually (Zeman and Cimprich, 2014).

We interpret the elevated γ-H2A.X levels on day 3 and 4 p.i. (Fig. 9A) to reflect the cells’ highly proliferative state in the pre-latent phase. This interpretation is clearly supported by the ATR inhibitor AZ20, which reduces the elevated γ-H2A.X levels in S and G2/M phase cells, only (Fig. 9D), whereas the ATM inhibitor KU-55933 is not effective and can be even counterproductive (Fig. 9C). In contrast to a previous publication (Nikitin et al., 2010) EBNA3A and EBNA3C do not seem to control a cellular DDR as indicated by measuring H2A.X phosphorylation (Fig. 9A) negating their roles in a presumed antiviral response of EBV infected cells. The slight increase of γ-H2A.X detection in cells infected with ΔEBNA3A/C EBV on day 5 and later (Fig. 9A) is in line with a moderately increased proliferation of these B cells (Fig. 8), but is inconsistent with the previous report by Nikitin et al. (Nikitin et al., 2010).

### The programmatic steps in the pre-latent phase

With the exception of B-lymphocytes infected with the ΔEBNA2 (5968) EBV mutant, all EBV infected cells show the same temporal pattern of cellular activation, cell cycle entry, and continuous cell proliferation during EBV’s pre-latent phase. This pattern defines clearly discernable phenotypes of the infected cells: (i) initial growth in cell size with a peak on day 4 p.i. (Fig. 2), (ii) first detection of DNA synthesis exactly on day 3 p.i. (Figs. 3–8, 10), (iii) occurrence of the first cell division on day 4 (ibid), (iv) a phase with very rapid cellular divisions until day 6 p.i. (ibid) that lead to (v) a decelerated, asynchronously proliferating cell populations, eventually. Only B cells infected with EBV mutants negative for LMP2A, EBNA-LP, EBNA1, or EBV’s many miRNAs showed a reduced rate of cell divisions in phase (iv), but the timing of all phases was not altered and extremely reproducible (Figs. 5-7, 10). This is surprising and speaks for a coordinated temporal reprograming of the infected cells that is intrinsic and probably depends on the early spike of EBNA2 expression on day 2 p.i. (Fig. 12A). It is attractive to speculate that MYC expression that follows suit is the only driver of this program (Kieffer-Kwon et al., 2017) until other viral factors contribute and stabilize the activated and reprogrammed cells. Interestingly, upon oncogenic induction of MYC (Macheret and Halazonetis, 2018) hyperproliferating cells have a short G1 phase. Its length is insufficient to correctly position and license replication origins leading to DRS in the subsequent S-phase (Macheret and Halazonetis, 2018). This recent finding is clearly in line with the phenotypes shown in Figure 9 supporting our conclusions.

The molecular events that initiate B cell activation seem to differ from those that lead to activated T cells upon antigen encounter or after CD3 and CD28 stimulation. This is because T lymphocytes react much more rapidly upon activation than B-lymphocytes and enter a hyperproliferative phase within 24 hours after activation (van Stipdonk et al., 2001; Slack et al., 2015 for a recent review). The reasons for this apparent difference are unknown and probably worth studying.

We took great care to find optimal conditions of B cell isolation and cultivation and choosing an optimal virus dose, which is a very critical parameter for the survival of EBV infected B cells (Fig. 1). However, independent of the source of the B-lymphocytes (secondary lymphatic tissue or PMBC) or their isolation procedure (FACS sorting or their “untouched” isolation by using magnetic beads-coupled depleting antibodies directed against surface antigens that B cell lack) we always noticed a considerable loss of viable cells during the first three days of infection accompanied by a variable but generally high fraction of Annexin V binding cells.

Currently, we do not know the origin of this observation. Given our optimal infection conditions it is very likely that EBV initially infects more than 90 % of the B-lymphocytes although the experiment shown in Fig. 1C detected considerably fewer EBNA2 positive cells on day 1 p.i.. This technical limitation is obvious when we analyze established lymphoblastoid cell lines by intracellular staining with our EBNA2 specific antibody. Depending on the individual cell sample this approach detects a fraction of about 60 to 90 % EBNA2 positive cells, only (data not shown), although all cells are expected to express this essential latent viral gene. EBER hybridization is the common method to detect latently EBV infected cells, but, in our hands, EBER hybridization was less sensitive compared with intracellular EBNA2 staining when the cells were analyzed shortly after infection.

As shown in Figure 1C higher MOIs resulted in larger fractions of EBNA2 positive cells on day 1 p.i. and slightly more activated cells compared with our standard MOI of 0.1 (Fig. 1D), but the initial loss of B-lymphocytes is comparable (Fig. 1A, B) and the number of B blasts on day 8 p.i. is reduced at higher MOIs at the start. The initial loss of B-lymphocytes is also apparent, when the cells are cultivated on CD40L expressing feeder cells in the presence of IL-4 (Fig. 4). A subpopulation of B-lymphocytes that is refractory to EBV or CD40L/IL-4 mediated activation and cell cycle entry would explain this initial failure of B cell survival. For example, certain B-lymphocytes might be incapable of becoming reprogrammed because their epigenetic or metabolic states prevent it. We have no evidence for this assumption but are currently investigating the different possibilities to identify and characterize the obstacles EBV encounters when it infects primary, resting B-lymphocytes.

## Materials and Methods

### Construction of mutant EBVs

All modifications of maxi-EBV plasmids described in this study rely on published techniques using homologous recombination in E.coli with linear DNA fragments (Warming et al., 2005). Following a recent development (Wang et al., 2009), we constructed a dual selection cassette consisting of the entire E.coli ribosomal S12 gene (*rpsL*) upstream of the aminoglycoside phosphotransferase gene (*aph*), which is also under the control of the *rpsL* promoter. Expression of *rpsL* results in streptomycin sensitivity at 1 mg/ml streptomycin sulfate in *rpsL*-deficient E.coli strains, whereas *aph* expression results in kanamycin resistance at 40 μg/ml kanamycin sulfate. This dual selection cassette was cloned onto the pJET1.2 (Fermentas) plasmid to yield the plasmid termed p6012. The entire *rpsL/kana* selection cassette is only 1348 bps in length and can be amplified with a PCR primer pair.

In this study all recombinant EBVs are all based on the maxi-EBV plasmid p2089, which comprises the entire B95.8 EBV genome cloned onto a mini-F-factor plasmid in E. coli (Delecluse et al., 1998). For this study, we modified the prokaryotic backbone of this maxi-EBV plasmid in E.coli to encode an artificial open reading frame consisting of the enhanced green fluorescence protein, eGFP, a T2A motif, and the gene encoding resistance against puromycin under the control of the human cytomegalovirus immediate-early promoter and enhancer. Puromycin resistance is conferred by the *pac* gene encoding a puromycin N-acetyl-transferase, and replaces the *hph* gene coding for hygromycin B phosphotransferase in p2089. The maxi-EBV plasmid based on p2089, which confers resistance against puromycin is called p6001 in our plasmid database (Tab. 1).

The EBV strain B98.8 suffers from a deletion that eliminates multiple miRNA loci, a second lytic origin of DNA replication, *oriLyt*, and additional genes that all field strains of EBV encode. We reconstructed a wild-type like strain based on B95.8 that expresses all viral miRNAs at their physiological levels. We introduced a DNA fragment derived from the EBV strain M-ABA and introduced it into the maxi-EBV plasmid p2089 to repair its deletion. In a second step, we replaced part of the bacterial mini-F factor backbone with the artificial open reading frame encoding both eGPF and *pac* as in p6001 described in the previous paragraph. The resulting maxi-EBV plasmid was very carefully analyzed with numerous restriction enzymes and DNA sequencing confirmed the two introduced loci and their intact flanking regions. The reconstructed wild-type EBV strain was termed r_wt/B95.8 (6008) (Tab. 1). It carries the right handed *oriLyt* and expresses all 25 EBV-encoded pri-miRNAs from their endogenous viral promoters as well as the LF1, LF2 and LF3 genes that B95.8 EBV lacks.

Subsequently, we replaced the viral miRNA loci with scrambled sequences as described for the previously published mutant EBV ΔmiR (4027) (Seto et al., 2010) abrogating the expression of all viral miRNAs. Extensive DNA sequencing confirmed all modified loci. The resulting maxi-EBV is called r_ΔmiR (6338) and is listed in Table 1. The complete sequence information of the 25 original and scrambled miRNA loci in r_ΔmiR (6338) and their secondary structure predictions can be found in Figure S7.

Based on p2089, its derivative p6001.1 or the newly constructed r_wt/B95.8 (6008) maxi-EBV we mutated viral loci or deleted them to obtain multiple EBV derivatives as listed in Table 1. In the following paragraphs we describe the individual experimental steps leading to functional knock-outs of EBNA3A, EBNA3C, and the concomitant knock-out of both viral genes. We also describe the construction of a pair of two maxi-EBVs, ΔEBNA-LP (5969) and its wild-type counterpart wt/B95.8 (5750). The construction of the other newly developed maxi-EBVs described in this study (Tab. 1) such as ΔEBNA2 (5968), ΔEBER (6431), ΔEBER/ΔmiR (6432), and ΔEBNA1 (6285) follows the same principles. Details of all EBV strains, their individual steps of construction and their annotated DNA sequence files are available upon request.

To establish a mutant EBV with a non-functional EBNA3C gene (Tab. 1), we modified p6001.1 by introducing the *rpsL/kana* cassette of p6012 into the EBNA3c locus. This insertional step was later reverted and replaced by a synthetic DNA fragment with two stop codons in the EBNA3C gene.

In detail, the *rpsL/kana* cassette in p6012 was PCR amplified with two primers, 5’-AGGGATGCTGCCTGCCGGGCTGTCAAGGTGAGTATGCCTCTAACTGGGTTCggcctggtgatgatggcgggatcg and 5’-ACCCTGTTAGGCACGGGAGTTAATGTGCGTAGTGTTGCTGTACGATATCCtcagaagaactcgtcaagaaggcg. The lower-case residues are homologous to sequences in p6012, the upper-case residues provide the homologous flanks of 50 nucleotides each, which direct the PCR fragment to recombine with the corresponding sequences in p6001.1, located at the 5’-end of EBNA3C from nucleotide coordinates #98,704 to #98,754 in the upper strand and #98,991 to #99,040 in the lower strand of the genomic B95.8 sequence, respectively. Recombination was achieved in the E.coli strain SW105 (a derivative of the DH10B strain and a kind gift from Neal G. Copeland, Mouse Cancer Genetics Program, National Cancer Institute, Frederick) harboring p6001.1 according to published protocols (Warming et al., 2005) to yield the maxi-EBV plasmid p6212.1, which was carefully analyzed by restriction enzyme cleavage and sequencing with primers confirming the correct insertion of the *rpsL/kana* containing PCR fragment.

Next, we used a synthetic DNA fragment (Genscript) of 712 bp in length termed p6122, covering nucleotide coordinates #98,505 to # 99,216 in B95.8. This EBV fragment encompasses parts of the first and second exons of EBNA3C and carries two TGA stop codons, which are in phase with the EBNA3C reading frame terminating translation after amino acids #144 in the first and #209 in the second exon of EBNA3C, corresponding to nucleotide coordinates #98,755 to #98,757 and #98,988 to #98,990 in B95.8 DNA, respectively. This DNA fragment released from p6122 with Asp718I cleavage was used to replace the *rpsL/kana* cassette in p6212.1 as described (Warming et al., 2005). In brief, recombination-competent SW105 cells were electroporated with this DNA fragment and subsequently selected on LB-plates with chloramphenicol (15 μg/ml) and streptomycin (1 mg/ml) at 32°C. The resulting maxi-EBV plasmid p6123.1, termed ΔEBNA3C (6123) throughout this manuscript, was carefully analyzed by cleavage with several restriction enzymes and partial DNA sequencing to confirm the two point mutations in EBNA3C and the correct insertion of the synthetic DNA fragment encompassing the two stop codons.

To inactivate the EBNA3A gene in p6123, we inserted the *rpsL/kana* gene cassette from p6012 into the EBNA3A start codon region in the maxi-EBV plasmid p6123. Two primer pairs, 5’-GGTACAAGGGGGGTGCGGTGTTGGTGAGTCACACTTTTGTTGCAGACAAAggcctggtgatgatggcgggatcg and 5’-AGGACGCCGAATTTTAGGGCGATGCCGAAAAGGTGTCAAGAAATATACAAcgatcccgccatcatcaccaggcc were used to amplify the *rpsL/kana* cassette (lower case nucleotide residues) and provide the flanking sequences for homologous recombination (upper case nucleotide residues) in p6123. The resulting PCR fragment was electroporated into the recombination-competent E. coli SW105 strain carrying the plasmid p6123. DNA recombination deleted 1348 bps of the first and part of the second exons of EBNA3A from nucleotide coordinates #92,243 to #93,590 in the genomic sequence of B95.8 using the *rpsL/kana* cassette as an insertional mutagen.

After selection on chloramphenicol/kanamycin plates, plasmid DNAs were prepared from *rpsL/kana* positive bacterial clones using the NucleoBond XtraBAC kit and carefully analyzed by restriction enzyme analysis. The resulting EBV plasmid termed ΔEBNA3A/C (6331) throughout this manuscript, was confirmed by DNA sequencing.

We generated a pair of recombinant EBVs with six copies of EBV’s BamHI-W-repeats (Fig. S2). The EBV recombinant wt/B95.8 (5750) is essentially wild-type (Tab.1), but its derivative ΔEBNA-LP (5969) is incapable of expressing EBNA-LP because each BamHI-W-repeat carries a translational stop codon in the W1 exon of EBNA-LP.

To construct the maxi-EBV plasmids pairs wt/B95.8 (p5750) and ΔEBNA-LP (p5969), we introduced the *galK* gene (Warming et al., 2005) into wt/B95.8 (p2089) deleting a large fragment of EBV DNA from nucleotide coordinate #12,001 to #47,634 removing the entire cluster of the BamHI-W-repeats. The resulting maxi-EBV plasmid was termed p5685.1. Next, we used a single copy of a BamHI-W-repeat. Its 5’ BamHI site was changed to a BglII site and a single nucleotide of its internal BglII site was mutated from 5’-AGATCT to 5’-GGATCT to inactivate it (Fig. S2B). This copy was multimerized to form a 6mer in pUC18 eliminating all internal BamHI or BglII sites. The both arms of the multimer was then equipped with EBV flanks to generate the plasmid p5537.1 in our database. The plasmid was cut with EcoRV and SphI to yield a DNA fragment of 24,896 bps. The left and right EBV flanks of this fragment with sequence homologies of 3005bps and 1716bps, respectively, mediated a successful DNA recombination with the recipient maxi-EBV plasmid p5685.1 to yield the final wt/B95.8 (5750) maxi-EBV plasmid (Tab. 1).

Similarly, for the construction of ΔEBNA-LP (p5969), we generated a 6-mer of a BamHI-W-repeat that is identical to the one in the previous paragraph but carries an additional mutation to inactivate the EBNA-LP gene. Figure S2B illustrates the situation at the sequence level of a single W1 exon. An XbaI site replaced the codons four to seven in the W1 exon introducing a translational stop signal at the forth codon (in the W1’ exon, the stop codon TAG replaces the second codon) abrogating the translation of any EBNA-LP transcript. The multimer was then equipped with EBV flanks to generate the plasmid p5571.2 in our database. This plasmid was cut with EcoRV and SphI to yield a DNA fragment of 24,896 bps as described in the paragraph above. DNA recombination with the recipient maxi-EBV plasmid p5685.1 led to the ΔEBNA-LP (p5685) maxi-EBV plasmid in our database. The backbone of this maxi-EBV plasmid was further altered to remove the *hph* gene but confer resistance against puromycin similar to wt/B95.8 (p6001) to yield the final ΔEBNA-LP (p5969) maxi-EBV plasmid (Tab. 1). The maxi-EBV plasmids wt/B95.8 (p5750) and ΔEBNA-LP (p5969) were very carefully scrutinized for any sequence alteration with the aid of multiple restriction enzymes and extensive DNA sequencing confirming the genetic compositions of the pair of maxi-EBV plasmids.

For the establishment of producer cell lines, DNAs of all maxi-EBV plasmids listed in Table 1 were further purified by two rounds of CsCl-ethidium bromide density gradient ultracentrifugations prior to their introduction into HEK293 cells.

### Cells and culture

All cells were maintained in RPMI 1640 medium supplemented with 10 % FCS, penicillin (100 U/ml), and streptomycin (100 μg/ml). Cells were cultivated at 37°C in a 5 % CO_2_ incubator.

### Preparation and quantification of infectious viral stocks

On the basis of HEK293 cells (obtained from the Leibniz Institut DSMZ - Deutsche Sammlung von Mikroorganismen und Zellkulturen GmbH, Braunschweig, Germany), virus producer cell lines were established after individual transfection of the maxi-EBV plasmid DNAs and subsequent selection with puromycin (500 to 1000 ng/ml). To obtain virus stocks, clonal producer cells were established on the basis of HEK293 cells, which transiently transfected with expression plasmids encoding BZLF1 and BALF4 to induce EBV’s lytic cycle. Three days post transfection, supernatants were harvested and centrifuged at 1200 rpm for 10 min and 4800 rpm for 10 min to remove cell debris. The titers of the different virus stocks were quantified and the concentrations of GFP-transducing virions expressed as ‘‘green Raji units’’ (GRUs) were determined as described recently in detail (Steinbrück et al., 2015). 1×10^5^ Raji cells (obtained from the Leibniz Institut DSMZ - Deutsche Sammlung von Mikroorganismen und Zellkulturen GmbH, Braunschweig, Germany) were incubated with different aliquots of virus stocks at 37°C for three days. The percentages of GFP-positive cells were determined by FACS using a BD FACS Canto instrument (BD Bioscience) and the linear regression equation was calculated as described (Steinbrück et al., 2015).

### Quantitation of B95.8 virus stocks

B95.8 virus supernatants were harvested from B95.8 cells (from the institute’s cell strain collection) and centrifuged to remove cellular debris. 2×10^5^ Elijah cells (Rowe et al., 1985) (from the institute’s cell strain collection) were incubated with defined volumes of B95.8 virus stock or a calibrated wt/B95.8 (2089) EBV stock, for which the concentration of GRU/ml was known. The cells were incubated at 4°C for 3 hours on a roller mixer, washed twice with PBS and the cell pellet was resuspended in 50 μl staining buffer and the anti gp350 antibody coupled to Alexa Fluor 647. After incubation at 4°C for 20 min, the cells were washed in staining buffer, resuspended in 300 μl staining buffer and the MFI of the stained cells were recorded. A linear regression equation was calculated on the basis of the applied amounts of wt/B95.8 (2089) EBV stock versus MFI, which allowed to deduce the concentration of the B95.8 virus stock.

### Isolation, separation, and infection of human primary B-lymphocytes

Peripheral B-lymphocytes were obtained from 500 ml peripheral blood as buffy coats with a starting volume of about 20 ml. The cells were diluted ten times with PBS, 0.5 mM acetylsalicylic acid, 1 mM EDTA to prevent platelet activation. The cells were purified by Ficoll density gradient centrifugation and the cell-containing interphase was washed several times (1600rpm, 1400rpm, 1200rpm for 10 min each). Remaining erythrocytes were removed with red blood lysis buffer (155 mM NH_4_Cl, 12 mM NaHCO_3_, 0.1 mM EDTA). 2 to 3×10^8^ PBMCs were taken up in 2.1 ml staining buffer (PBS, 0.5 % BSA, 2 mM EDTA) and 300 μl of anti-human CD61 and CD3 microbeads (Miltenyi Biotec) were added to remove contaminating platelets and T cells, respectively, on four LD columns by magnetic sorting according to the manufacturer’s protocol (Miltenyi Biotec).

Human primary B cells from adenoids were separated from T cells by rosetting with sheep erythrocytes (purchased from Thermo Scientific Oxoid, cat. no. SR0051D) and purified by Ficoll density gradient centrifugation as described in the previous paragraph. Naïve B-lymphocytes (IgD^+^/IgH^+^, CD38^-^, CD27^-^) were physically sorted on an Aria III instrument (Becton Dickinson) using the following antibodies: CD38 (eBioscience, #25-0389-42), IgD (BD Pharmingen, #555778), CD27 (BD Pharmingen, #337169). For virus infection, primary B cells obtained from anonymous donors (see Ethics Statement below) were incubated with virus stocks at a multiplicity of infection of 0.1 GRU for 20 hours. After replacement with fresh medium, the infected cells were seeded at an initial density of 1×10^6^ cells per ml in 96 or 48-well cluster plates.

B-blasts were generated after plating sorted naïve B-lymphocytes on irradiated CD40 ligand feeder cells in the presence of IL-4 at 2 ng/ml as described (Wiesner et al., 2008).

### Ethics Statement

PBMCs in the form of buffy coats from healthy adult donors were purchased from the Institute for Transfusion Medicine, University of Ulm, Germany. The buffy coats were a side product of standard collection of erythrocytes for medical blood transfusion and were delivered to us in fully anonymized form.

Where indicated, primary naïve B-lymphocytes were isolated from adenoid samples. They were leftovers from adenoidectomies performed at the Department of Otorhinolaryngology and Head and Neck Surgery, Klinikum der Universität München; these samples also were transferred to our institution in fully anonymized form. The institutional review board (Ethikkommission, Klinikum der Universität München, Munich, Germany) approved this procedure (project no. 071-06 - 075-06).

### Antibodies for FACS analyses or Western blot immunodetection

For intracellular FACS staining, an Alexa Fluor 647 conjugated mouse monoclonal antibody directed against phosphorylated H2A.X (γ-H2A.X) (clone N1-413) was purchased from BD Bioscience (Cat. No. 560447). The rat anti-EBNA2 (1E6) antibody was conjugated with Alexa Fluor 647 and obtained from the in-house Monoclonal Antibody Core Facility.

For Western Blotting, the primary mouse monoclonal antibody against Ku70 (clone A-9) was purchased from Santa Cruz, the rabbit monoclonal antibody specific for p21 (clone EPR3993) was purchased from Abcam. Polyclonal rabbit antibodies directed against p53 (#9282) and Rad51 were purchased from Cell Signaling and Santa Cruz, respectively. Rat antibodies specific for EBNA2 (R3 and 1E6), EBNA1 (1H4), and EBNA3A (clone E3AN 4A5-11111) proteins and mouse antibodies specific for EBNA-LP (JF186), MYC (9E10), and EBNA3C proteins (clone E3A10-P2-583) obtained from the in-house Monoclonal Antibody Core Facility were used.

### Intracellular FACS staining of **_γ_**-H2A.X

The infected cells were treated with 85 μM etoposide for 1 h or left untreated before harvest at the indicated times points p.i.. Intracellular FACS stainings were performed with fixed cells using the Fix and Perm Kit (Thermo Fisher Scientific). Briefly, 5×10^5^ cells were washed and suspended in 100 μl PBS. Cells were fixed by adding 100 μl Fix and Perm Medium A for 15 min in the dark at room temperature. The fixed cells were washed with 1 ml staining buffer (PBS, 0.5 % BSA, 2 mM EDTA) and the pellet resuspended in 1 ml freezing medium (90% FCS, 10% DMSO) to be stored at −80°C. After sample collection was completed, the cells were thawed, washed again, centrifuged and the pellets were resuspended and permeabilized in 100 μl Fix and Perm Medium B. During the permeabilization step, anti γ-H2A.X staining was performed for 20 min at room temperature in the dark. γ-H2A.X specific mouse monoclonal antibody conjugated Alexa Fluor 647 were diluted 1:50. After another washing step with 1 ml staining buffer the samples were measured with a BD Fortessa or Canto Cell Analyzer (Becton Dickinson).

### Intracellular FACS staining of EBNA2

The infected cells were harvested at the given time points after infection, pelleted by centrifugation and resuspended in 100 μl PBS. Cells were fixed by adding 100 μl Fix and Perm Medium A (Life Technologies) for 15min in the dark at room temperature. The fixed cells were washed with 1 ml staining buffer (PBS, 0.5% BSA, 2mM EDTA), pelleted and resuspended in 1 ml freezing medium (90% FCS, 10% DMSO) to be stored at −80°C. After sample collection, the cells were thawed and washed in 1 ml staining buffer. The pellet was resuspended in 100 µl Fix and Perm Medium B (Life Technologies) and 1.7 µl of a 1:10 dilution of the anti-EBNA2 monoclonal antibody 1E6 coupled with Alexa Fluor 647 (1:600 final) was added. The cells were incubated at room temperature for 20 min, washed and resuspended in staining buffer for FACS analysis.

### BrdU incorporation and cell cycle analysis

Metabolic BrdU labeling and detection of newly synthesized DNA were performed by adding 10 µM BrdU (APC BrdU Flow Kit, BD Pharmingen, # 557892) for one hour prior to harvest. The cells were washed in staining butter (PBS, 0.5% BSA, 2mM EDTA) and the pellets were resuspended and fixed in 100 μl Cytofix/Cytoperm buffer (APC BrdU Flow Kit) and incubated on ice for 20 min. The cells were washed in staining buffer, pelleted and resuspended in 1 ml freezing medium (90% FCS, 10% DMSO) to be stored at −80°C. After sample collection, the cells were thawed and washed in 1 ml staining buffer. The pellet was resuspended in 100 µl Cytofix/Cytoperm buffer (APC BrdU Flow Kit) incubated on ice for 5 min, washed in 1 ml Perm/Wash buffer (APC BrdU Flow Kit), and further processed according to the manufacturer’s instructions. Cellular DNA was counterstained with 7-AAD (BioLegend) and the cells were analyzed on a BD LSR Fortessa instrument.

### EdU incorporation and inhibitors of DDR and DRS

Metabolic EdU labeling and detection of newly synthesized DNA was performed with the Click-iT EdU flow cytometry cell proliferation assay by Thermo Fisher Scientific and Alexa Fluor 488 azide. The cells were metabolically labeled with 10 µM EdU for one hour and analyzed following the manufacturer’s instructions. KU-55933 (SML 1109) and AZ20 (SML 1328) were purchased from Sigma-Aldrich and used at final concentrations of 10 µM and 2 µM, respectively.

### Cell size, TMRE staining

Primary naïve B-lymphocytes were infected with wt/B95.8 (2089) EBV with an MOI of 0.1 and analyzed for their cell diameter daily. Briefly, we purified intact cells by Ficoll density gradient centrifugation, washed the cells in PBS and recorded microscopic images of the cell samples using a Tali image-based cytometer (Thermo Fischer). We interpreted the images using the ImageJ software package and identified cellular objects with circularity >0.75 and aspect ratio ≦1.3 as intact cells. The individual diameters of each cell were calculated based on images obtained with calibration beads of a radius of 1.9 mm on average (Tali Calibration Beads; Thermo Fischer). Mean and standard deviation of the cells’ diameters were determined accordingly (GraphPad Prism software, vers. 7) and the volumes of cells were calculated assuming a spherical shape. The incorporation of TMRE has been described recently (Mrozek-Gorska et al., 2019).

### Protein lysates and Western blot immunodetection

The infected cells were harvested at the indicated times post-infection. Cells were washed with phosphate-buffered saline (PBS), and lysed with lysis buffer (0.02 % sodium dodecyl sulfate [SDS], 0.5 % Triton X-100, 300 mM NaCl, 20 mM Tris-HCl [pH 7.6], 1 mM EDTA, 1 mM dithiothreitol) for 10 min on ice. Samples were centrifuged at 10,000 g for 1 min at 4°C. Samples were adjusted to 5×10^7^ cells/ml lysis buffer.

6 % polyacrylamide gels were used for EBNA3A and EBNA3C, for other proteins, 12 % polyacrylamide gels were used. 10 μl of protein samples (lysate from 5×10^5^ cells) were loaded per lane. Protein samples were separated by SDS-PAGE and subsequently transferred onto 0.45 μM nitrocellulose membranes (Bio-Rad) for 60 min at 100 V in a Mini-PROTEAN Tetra Cell (Bio-Rad). For EBNA3A and EBNA3C detections, proteins were transferred for 90 min. Membranes were blocked with 5 % milk in PBS containing 0.05 % Tween-20 (PBS-T) for 1 h at room temperature. Membranes were probed as indicated; the anti-Ku 70 mouse monoclonal antibody was diluted 1:1,000 and incubated for 1 h at room temperature; the anti-p21 rabbit monoclonal antibody was diluted 1:5,000 and incubated at 4°C overnight; the anti-p53 rabbit polyclonal antibody was diluted 1:1,000 and incubated at 4°C overnight; with anti-Rad51 rabbit polyclonal antibody was diluted 1:500 and incubated at 4°C overnight; and the anti-EBNA2 rat polyclonal antibody was diluted 1:200 and incubated at 4°C overnight. Horseradish peroxidase (HRP)-conjugated species-specific goat secondary antibodies were used at a 1:5,000 dilution in PBS-T and the membranes were incubated for 1 h at room temperature. The signals were visualized by exposure to X-ray film.

## Acknowledgements

Acknowledgements and Funding

We thank Christine Göbel, Munich, for her precious experimental support and expertise and Takanobu Tagawa, Boston, for computing the cell sizes from digital images. This work was financially supported by grants of the Deutsche Forschungsgemeinschaft [grant numbers SFB1064/TP A13, SFB-TR36/TP A04], Deutsche Krebshilfe [grant number 70112875], and National Cancer Institute [grant number CA70723] to W.H.. A.S. was supported by a ‘Humboldt Research Fellowship’ and by two postdoctoral grants from ‘The Uehara Memorial Foundation’ and an ‘JSPS Research Fellowship’. M.B. received an EMBO long-term fellowship by the European Molecular Biology Organization.

## Authors contributions

D.P., P.M.-G., M.B., A.S., E.A., and W.H. designed and performed experiments and analyzed data. A.G., S.H., P.L., and W.H. designed and conceived the project. W.H. and S.H. wrote the manuscript.

## Declaration of Interests

The authors declare no competing interests.

