## Supplemental Text and Figures S1-S7 for "The first days in the life of naïve human B-lymphocytes infected with Epstein-Barr virus"

### Supplemental Information

Dagmar Pich[1], Paulina Mrozek-Gorska[1], Mickaël Bouvet[1], Atsuko Sugimoto[1†], Ezgi Akidil[1], Adam Grundhoff[2], Stephan Hamperl[3], Paul D. Ling[4], Wolfgang Hammerschmidt[1*]

*Corresponding author

[1]Research Unit Gene Vectors, Helmholtz Zentrum München, German Research Center for Environmental Health and
German Center for Infection Research (DZIF), Partner site Munich
Marchioninistr. 25
D-81377 Munich, Germany

[2]Heinrich Pette Institute, Leibniz Institute for Experimental Virology
Martinistr. 52
D-20251 Hamburg, Germany

[3]Institute of Epigenetics and Stem Cells, Helmholtz Zentrum München, German Research Center for Environmental Health
Marchioninistr. 25
D-81377 Munich, Germany

[4]Department of Molecular Virology and Microbiology
Baylor College of Medicine
Houston, Texas, United States

[†]current address: Kyoto Pharmaceutical University, Department of Cell Biology, Misasagi-Shichonocho 1, Yamashinaku, Kyoto-shi, Kyoto 607-8412, Japan

### Legends of Supplemental Figures

Fig. S1. Activation kinetics of peripheral B-lymphocytes isolated from PBMC and infected with wt/B95.8 (2089) EBV.

Resting B-lymphocytes from PBMCs were infected with wt/B95.8 (2089) EBV with an MOI of 0.1 and analyzed for the division index, cell numbers, and percentage of Annexin V-positive cells. Infection experiments with PBMCs from three different donors were performed, shown is one representative set of results.

Fig. S2. Genetic map of the EBNA-LP and EBNA2 loci of wt/B95.8 (5970) and ΔEBNA-LP (5969) EBV.

A. Details of the two EBV strains wt/B95.8 (5970) and ΔEBNA-LP (5969) engineered to carry six copies of the BamHI-W-repeats are shown. Indicated are the two cis-acting elements *oriP* (with its family of repeats and dyad symmetry elements, FR and DS, respectively) and *oriLyt* and the exons encoding the EBNA-LP gene (W0, $[W1, W2]_6$, Y1, Y2), EBNA2, BHLF1, and BHRF1. The BamHI-W-repeats are flanked by XhoI sites, the BamHI sites preserved in wt/B95.8 (5970) and ΔEBNA-LP (5969) are indicated. Two alternative splicing forms of the bicistronic EBNA-LP/EBNA2 transcripts initiating from either the Cp or Wp promoter are shown below the genetic maps.

B. The schematic composition of the first BamHI-W-repeat with parts of its preceding BamHI-C fragment in two EBV strains is shown together with the relevant exons C2, W0, W1/W1', and W2. The restriction enzyme sites BamHI and BglII are indicated in the EBV strain wt/B95.8 (5970) that are altered in ΔEBNA-LP (5969). In ΔEBNA-LP (5969) each copy of the BamHI-W-repeat carries a translational stop codon in the W1 exon indicated by an XbaI site terminating the translation of the EBNA-LP gene. The codon usage in the W1 exon of the wt/B95.8 (5970) and ΔEBNA-LP (5969) EBV strains is provided.

Fig. S3. Steady-state levels of EBNA2 and EBNA-LP proteins in B cells infected with three different EBV strains.

Naïve B-lymphocytes were isolated from adenoid tissue of two different donors and infected with wt/B95.8 (2089), wt/B95.8 (5750), or ΔEBNA-LP (5969) EBV at an MOI of 0.1. Cells were cultivated for seven (experiment A) or eight weeks (experiment B) and protein extracts from B cells were analyzed with antibodies specific for EBNA2 or EBNA-LP as indicated. ΔEBNA-LP (5969) EBV infected cells did not express EBNA-LP as expected. Two experiments out of three are shown.

Fig. S4. Analysis of cell proliferation and Annexin V binding of B cells infected with mutant EBVs negative for viral non-coding RNAs.

Naïve B-lymphocytes were isolated from adenoid tissue, physically sorted and infected with four different EBV strains as indicated. Their genotypes are summarized in Table 1. The cell numbers and the fraction of Annexin V-positive cells were analyzed daily. One representative experiment out of four is shown.

Fig. S5. Western blotting analysis of proteins regulated during cellular DNA damage response.

Uninfected human primary B-lymphocytes (uninf.) and cells infected with wt/B95.8 (2089) EBV or ΔEBNA3A/C (6331) EBV were harvested at the indicated time points (days p.i.). Protein lysates of $5\times10^5$ cells per lane were loaded and the steady state levels of the indicated proteins were analyzed with antibodies directed against p53, p21, Ku70, or Rad51. An EBNA2 specific antibody was used to monitor the onset of EBNA2 expression. Lysates obtained from 293T cell incubated with 85µM etoposide for one hour were loaded as control (cont). One experiment out of two is shown.

Fig. S6. FACS based cell size analysis of B-lymphocytes infected with ΔEBNA1 (6285) or wt/B95.8 (2089).

Human primary B cells from adenoid were infected with wt/B95.8 (2089) or ΔEBNA1 (6285) EBV with an MOIs of 0.1 and analyzed by flow cytometric analysis according to forward (FSC-A; x-axis) and sideward (SSC-A; y-axis) scatter criteria at the indicated time points. Viable cells are surrounded by the indicate gate (polygonal red line). Cells infected with the two EBV strains similarly gain in size and granularity until eight days p.i., but B cells infected with ΔEBNA1 (6285) EBV showed a decrease in volume starting on day 10 p.i. and the main population of cells became very small two weeks p.i. compared with cells infected with wt/B95.8 (2089) EBV. Shown is one representative experiment out of three.

Fig. S7. Alignments and predicted structures of mutant miRNAs.

This multi-page figure shows alignments (pages 1 to 5) and predicted secondary structure images (pages 6 to 12) of the 25 pre-miRNAs encoded by EBV field strains (represented by the EBV GenBank entry AJ507799) and the corresponding sequences in the mutant EBV strain r_ΔmiR (6338) that carries 25 scrambled miRNA genes. Secondary structures were predicted using the Vienna RNAfold package [1] and are indicated by bracket notation above and below the aligned sequences. Regions that encode mature miRNA sequences or their scrambled counterparts are shown in boldface on grey background in the alignments, or in red in the structure images. The labelling to the left of the aligned sequences denotes the particular miRNAs that correspond to the AJ507799 sequence (shown at the top of each alignment and structure image pair), or is mutated in the r_ΔmiR (6338) EBV strain. The region encoding the mature ebv-miR-BART4* and ebv-miR-BART5* were not scrambled in r_ΔmiR (6338), but the mutations of the mature ebv-miR-BART4 and ebv-miR-BART5 sequences efficiently destroy the pre-miRNA hairpin structures (see structure prediction on pages 7 and 8 for these two miR sequences) ablating the expression of ebv-miR-BART4* and ebv-miR-BART5*.

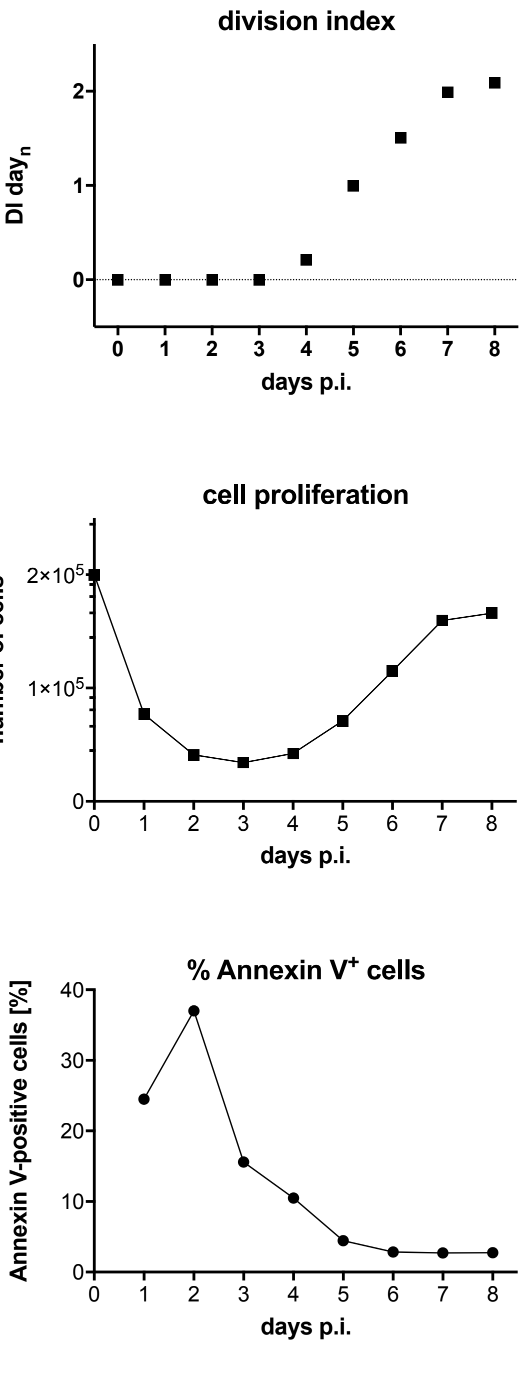

division index
DI $day_n$
2
1
0
0 1 2 3 4 5 6 7 8
days p.i.
cell proliferation
number of cells
$2\times10^5$
$1\times10^5$
0
0 1 2 3 4 5 6 7 8
days p.i.
% Annexin V+ cells
Annexin V-positive cells [%]
40
30
20
10
0
0 1 2 3 4 5 6 7 8
days p.i.


Fig. S1

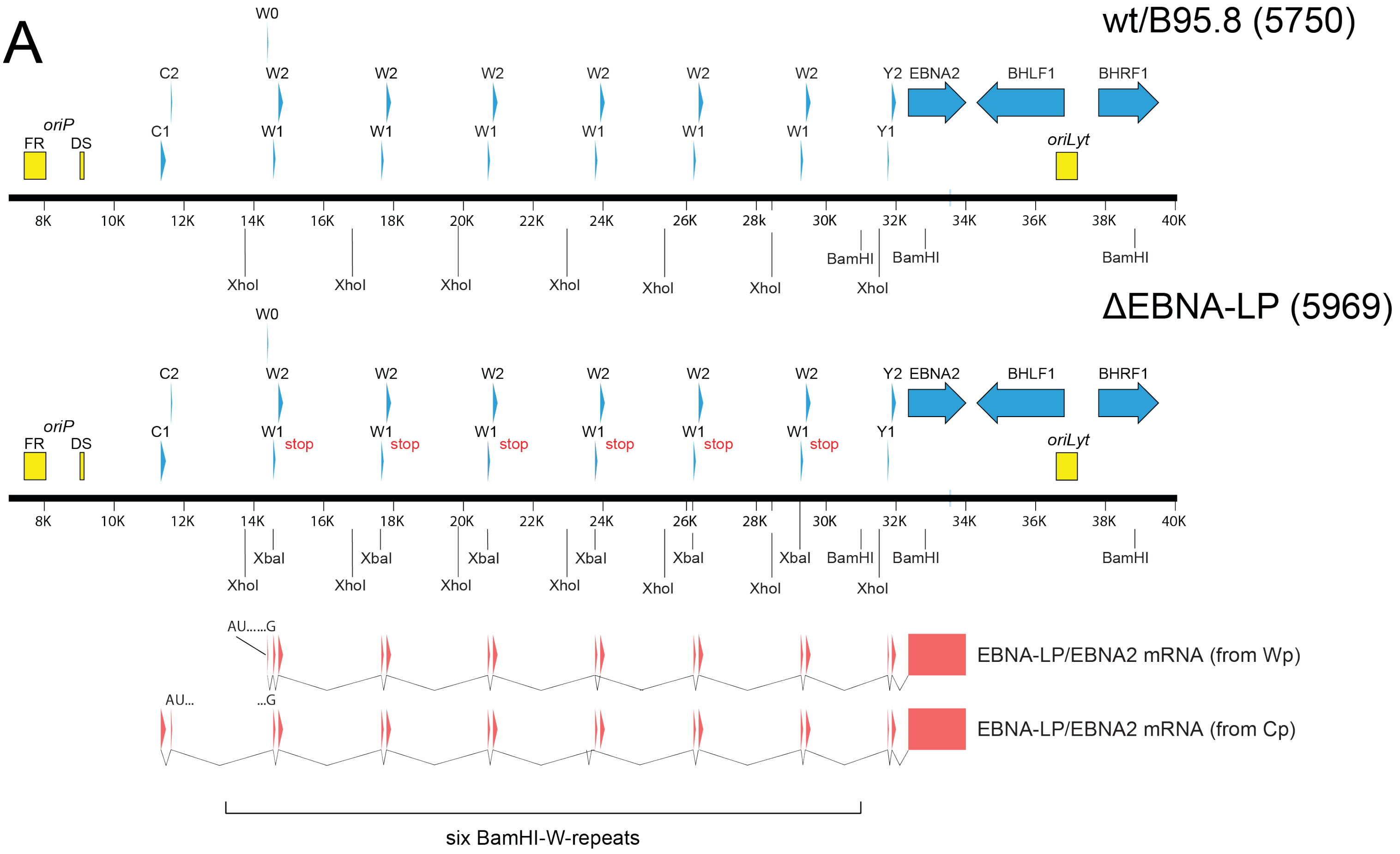


B

C2/BamHI-W-repeat array in wt/B95.8 (2089)

C2------BamHI------BglII-------W0---W1/W1'---W2------BamHI---

```
          GGATCC        AGATCT                                       GGATCC
          CCTAGG        TCTAGA                                       CCTAGG
```

C2/BamHI-W-repeat array in wt/B95.8 (5750) and ΔEBNA-LP (5969)

C2-----(BamHI)----(BglII)------W0---W1/W1'---W2-----(BamHI)--

```
          GGATCT        GGATCT                                      GGATCT
          CCTAGA        CCTAGA                                      CCTAGA
```

W1 exon in wt/B95.8 (5750)

```
CCT AGG GGA GAC CGA AGT GAA GGC CCT GGA CCA ACC CGG CCC GGG CCC CCC GGT ATC GGG CCA GAG
GGA TCC CCT CTG GCT TCA CTT CCG GGA CCT GGT TGG GCC GGG CCC GGG GGG CCA TAG CCC GGT CTC
 P   R   G   D   R   S   E   G   P   G   P   T   R   P   G   P   P   G   I   G   P   E
```

W1 exon in ΔEBNA-LP (5969)

```
CCT AGG GGA TAG TCT AGA CTA GGC CCT GGA CCA ACC CGG CCC GGG CCC CCC GGT ATC GGG CCA GAG
GGA TCC CCT ATC AGA TCT GAT CCG GGA CCT GGT TGG GCC GGG CCC GGG GGG CCA TAG CCC GGT CTC
 P   R   G   *
```

Fig. S2

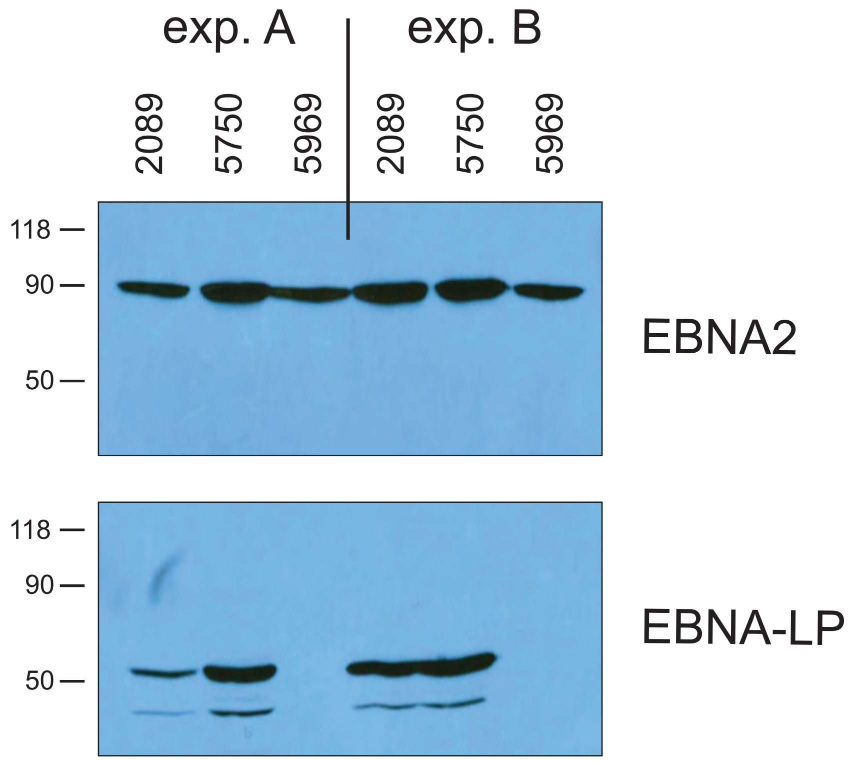
exp. A
exp. B
2089
5750
5969
2089
5750
5969
118
90
50
EBNA2
118
90
50
EBNA-LP

Fig. S3

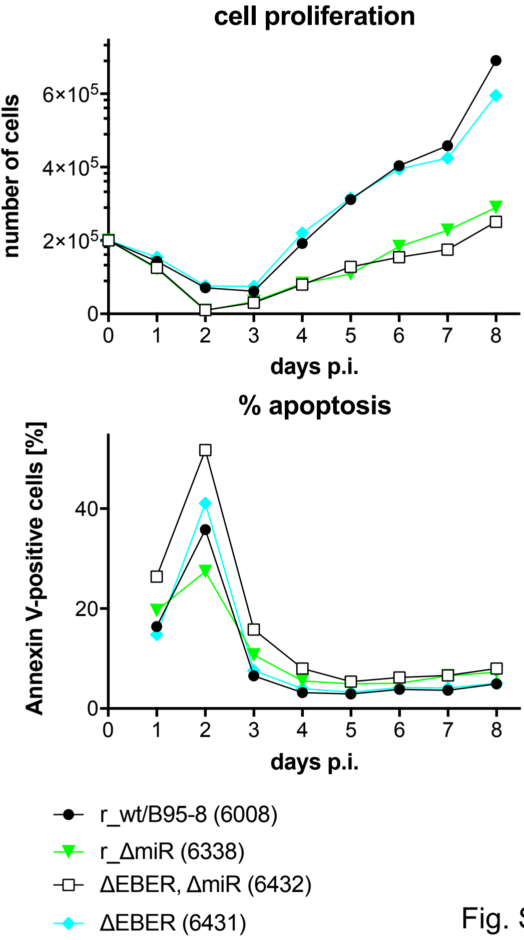
cell proliferation
number of cells
6×10^5
4×10^5
2×10^5
0
0
1
2
3
4
5
6
7
8
days p.i.
% apoptosis
Annexin V-positive cells [%]
40
20
0
0
1
2
3
4
5
6
7
8
days p.i.
r_wt/B95-8 (6008)
r_ΔmiR (6338)
ΔEBER, ΔmiR (6432)
ΔEBER (6431)

Fig. S4

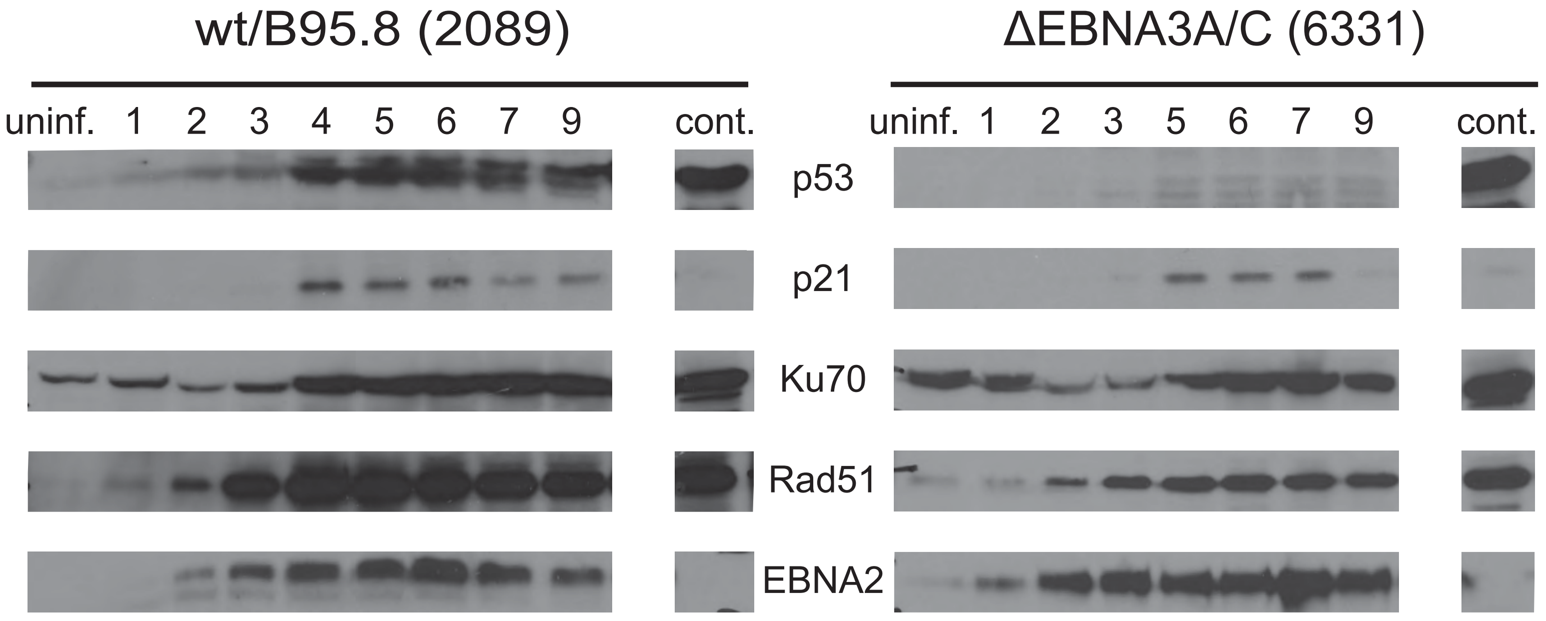
wt/B95.8 (2089)
ΔEBNA3A/C (6331)
uninf. 1 2 3 4 5 6 7 9 cont.
uninf. 1 2 3 5 6 7 9 cont.
p53
p21
Ku70
Rad51
EBNA2

Fig. S5

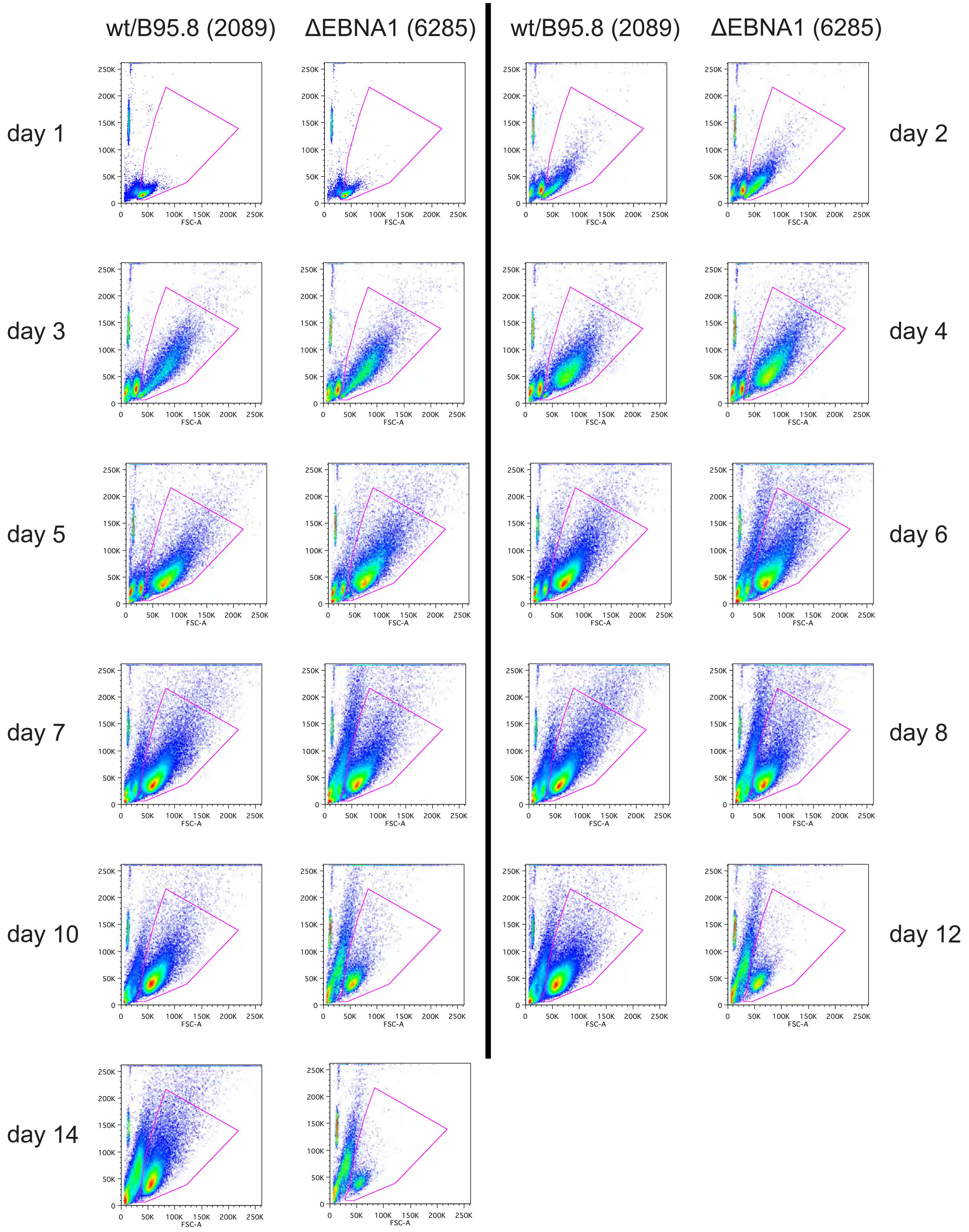

wt/B95.8 (2089)
ΔEBNA1 (6285)
wt/B95.8 (2089)
ΔEBNA1 (6285)
day 1
day 2
day 3
day 4
day 5
day 6
day 7
day 8
day 10
day 12
day 14
0
50K
100K
150K
200K
250K
FSC-A


Fig. S6

ebv-miR-BHRF1-1

```
                    ebv-miR-BHRF1-1
              (((.((((.(((((((..((((.(((((.((((.............)))).))))).))))..)))))))....)))))))
AJ507799      GGCUCCUUAUUAACCUGAUCAGCCCCGGAGUUGCCUGUUUCAUCACUAACCCCGGGCCUGAAGAGGUUGACAAGAAGGGUC
              |||||||||| |   | | | |        |||||||||||||||||||||||||||||||||||||||||||||||||
r_ΔmiR(6338)  GGCUCCUUAUGAUAAUAACCCGGGUGCCCCUUGCCUGUUUCAUCACUAACCCCGGGCCUGAAGAGGUUGACAAGAAGGGUC
              (((.((((.((.......((((((.((.....))..(((........))))))))(((....))....))..)))))))
```

ebv-miR-BHRF1-2

```
                      ebv-miR-BHRF1-2*                    ebv-miR-BHRF1-2
              ((((..(((((((((.(((((((((((.(((((((...........))))))).))))))))))).)))))))))..))))
AJ507799      GGCCCCCACUUUUAAAUUCUGUUGCAGCAGAUAGCUGAUACCCAAUGUUAUCUUUUGCGGCAGAAAUUGAAAGUGCUGGCC
              |||||||||||||    |           || |   ||||||||||||| |   ||    |    |  |||  |||||||||||
r_ΔmiR(6338)  GGCCCCCACUUUUCGGUAGGUGAUAUCCAAACUAUUGAUACCCAAUGUGAAACUUCUGGAUCGUGAUUAUAAGUGCUGGCC
              ((((..(((((.(((((((............)))))))...(((............)))...........)))))..))))
```

ebv-miR-BHRF1-3

```
                      ebv-miR-BHRF1-3
              ((((((.((((.((((((..((((((.((((((...(((.........))))))))))))))).))))))).)))).))).)))
AJ507799      CCUGUGGUGUUCUAACGGGAAGUGUGUAAGCACACACGUAAUUUGCAAGCGGUGCUUCACGCUCUUCGUUAAAAUAACACAAGG
              ||||||||||||| | |  |||       | |  |||||||||||||||||||||||||||||||||||||||||||||||||
r_ΔmiR(6338)  CCUGUGGUGUUCUGAAGAUAAGGUCCACGGAAUGCACGUAAUUUGCAAGCGGUGCUUCACGCUCUUCGUUAAAAUAACACAAGG
              ((((((((((((((..(((...)))..)))))))))............(((((.((.....))...)))))........))).)))
```

ebv-miR-BART3

```
                    ebv-miR-BART3*                       ebv-miR-BART3
              (((((((((..((((.(((((((((.(((((((............))))))).)))))).))).))))..)))))))))
AJ507799      CCUUUGGUGGAACCUAGUGUUAGUGUUGUGCUGUAAAUAAGUGUCCAGCGCACCACUAGUCACCAGGUGUCACCGGAGG
              ||||||||||| |||||| |  ||  |||  ||||||||||||||||| |   |         | | |  | |||||||||
r_ΔmiR(6338)  CCUUUGGUGGAGCCUAGUUUCGGUUGUGUAAUGUAAAUAAGUGUCCAGUGUGUCCACCACAAACUGCCGGCACCGGAGG
              (((((((((.((((.......))))...................((((.(((.....))).))))....)))))))))
```

ebv-miR-BART4

```
                      ebv-miR-BART4                           ebv-miR-BART4*
              ((...((((((((.(((((.((((((((((((((((............))))))))).)))).)).))))).))))))))..))
AJ507799      UAACAUUUGGUGGGACCUGAUGCUGCUGGUGUGCUGUAAAUAAGUGCCUAGCACAUCACGUAGGCACCAGGUGUCACCAGGGCUA
              ||||||||||||| |      |  ||      ||  |||||||||||||||||||||||||||||||||||||||||||||||||
r_ΔmiR(6338)  UAACAUUUGGUGGCAGAGCCUUUUGGGUCGGUCGUGUAAAUAAGUGCCUAGCACAUCACGUAGGCACCAGGUGUCACCAGGGCUA
              .....((((((((((((.((....)).))..............((((((.((.......)))))))....))))))))))....
```

Figure S7

```
ebv-miR-BART1
                    ebv-miR-BART1-5p                      ebv-miR-BART1-3p
              ((((.((.(((...(((((((((((..((((((((.........))))))))))).))))))))...))).))))))
AJ507799      CCCGUGGGGGUCUUAGUGGAAGUGACGUGCUGUGAAUACAGGUCCAUAGCACCGCUAUCCACUAUGUCUCGCCCGGG
              |||||||||||    |     | |  | |||    ||||||||||||| |        |  ||        |||||||||
r_ΔmiR(6338)  CCCGUGGGGGUGGGACCUUAUGGUAUGUGGAUCGAAUACAGGUCCACAAUUAGAUUCACCCUCCCUGGUCGCCCGGG
              .(((.(((((((((.(((....))).((((((((.......))))))))).......))))))))).))).........
```

```
ebv-miR-BART15
                                                          ebv-miR-BART15
              .(((...(((..(((((((((.((((((.(((..((((.....)))))))))))))).)))))))))....))).))).
AJ507799      UGUGCCGCUUGGAGGGAAACAUGACCACCUGAAGUCUGUUAACCAGGUCAGUGGUUUUGUUUCCUUGAUAGAGACACA
              ||||||||||||||||||||||||||||||||||||||||||||||     | ||||||||        ||||||||||
r_ΔmiR(6338)  UGUGCCGCUUGGAGGGAAACAUGACCACCUGAAGUCUGUUAACCAGUCGCUUUGUUUUGUUGUGACAGUAGAGACACA
              .(((...(((...((..((((.(((........)))))))..)).(((((...........)))))....))).))).
```

```
ebv-miR-BART5
                  ebv-miR-BART5                              ebv-miR-BART5*
              (((.(((....((((((((..(((.((((((((................)))))))).)))))))))))...))).)))
AJ507799      GUGGCACCUCAAGGUGAAUAUAGCUGCCCAUCGACGUAUCGCUGGAAACCGGUGGGCCGCUGUUCACCUAAAGUGACGC
              ||||||||||||||||||||||||||||||||||||||||||||||||||||||||||    |          |  |  |
r_ΔmiR(6338)  GUGGCACCUCAAGGUGAAUAUAGCUGCCCAUCGACGUAUCGCUGGAAACCGGUGGGCCAGGGCCUGGCCGCACGGGCCA
              ....((((....))))..........((((((((..((.((....)).)))))))))).((((.....))))........
```

```
ebv-miR-BART16
                       ebv-miR-BART16
              (((.(((((.((...((((((..(((((.(((.(((((...........))))).))))))))))))))...)).))))).)))
AJ507799      CAGGUGUGGAAUUUAGAUAGAGUGGGUGUGUGCUCUUGUUUAAUUACACCAAGAUCACCACCCUCUAUCCAUAUCCCACAAUUG
              ||||||||||||      |      |     ||||   ||||||||||||||||||||||||||||||||||||||||||||||||
r_ΔmiR(6338)  CAGGUGUGGAAUCGGAUUGAUACGAUGUUGUGGGUUUGUUUAAUUACACCAAGAUCACCACCCUCUAUCCAUAUCCCACAAUUG
              (((.(((((.((((........))))...((((.((((...........))))....))))...............))))).)))
```

```
ebv-miR-BART17
                 ebv-miR-BART17-5p                      ebv-miR-BART17-3p
              (((...(((((((.((((((((((((((((((............))))))))))).)))))).))))).))..)))
AJ507799      GUGGCACCCUAAGAGGACGCAGGCAUACAAGGUUAUUACCCAGUCCUUGUAUGCCUGGUGUCCCCUUAGUGGGACGC
              |||||||||     | |              | |||||||||||||||||| ||  || |        |      |||||||
r_ΔmiR(6338)  GUGGCACCCCCGAGGAAAAACUAACGGUAGGGUUAUUACCCAGUCCUCGUCAGCGUUCGUAUUUCGGUUCGGGACGC
              ..(((.....((((((...((.....)).((((....))))..))))))...((((((.............))))))
```

Figure S7 continued

```
ebv-miR-BART6
                         ebv-miR-BART6-5p                   ebv-miR-BART6-3p
               ((((....((((((((((((((((.((((...((.((((......)))).))..)))).))))))))))))))))))...))))
AJ507799       CUUGUUGGUACUUUAAGGUUGGUCCAAUCCAUAGGCUUUUUUUGUGAAAACCCGGGGAUCGGACUAGCCUUAGAGUAACUCAAG
               |||||||||||||||  |  | |    |        |||||||||||||||||  |    |              ||||||||||
r_ΔmiR(6338)   CUUGUUGGUACUUUCUGAAUCGCGAGAGUUUGCAACUUUUUUUGUGAAAACCAAGACCGCCACUGGUGUGAGUGGUAACUCAAG
               ((((.((((.(((((........)))))((..(((......)))..)).))))..((((((((....))).)))))....))))
```

```
ebv-miR-BART21
                         ebv-miR-BART21-5p                  ebv-miR-BART21-3p
               ((.((((((((((..((((((((...(((((((.(((........))).)))))))...))))))))..))))))).)))).))
AJ507799       GUAUGGGCUGGGUAUUCACUAGUGAAGGCAACUAACACAGUUAGACGUGCUAGUUGUGCCCACUGGUGUUUAUCCGGUCCCAAAU
               ||||||||||||     |    ||||  | ||  |   ||||||||    |       |   |   |  |   | ||||||||||
r_ΔmiR(6338)   GUAUGGGCUGGGGCCCAAGAGAUGAAAUCUACAUAUCUAGUUAGACUGCUUUUGGUGACGUUCCCAUAGUCUGCGGGUCCCAAAU
               .....((((((((((.(((.(((.......))).)))...((((((....((....))......))))))).)))))))))...
```

```
ebv-miR-BART18
                         ebv-miR-BART18-5p                  ebv-miR-BART18-3p
               ((((((..((((...(((((((.((((((.((((((.((....)).)))))).)))))).))))))).....)))))..))))))
AJ507799       CGGGUGUCCUGGCUCAAGUUCGCACUUCCUAUACAGUGUUAAAGCCUUGUAUCGGAAGUUUGGGCUUCGUCCCAGUGUACUCG
               ||||||||||||||| | |||       ||     ||||||||||||||     | ||     | | |   ||||||||||||
r_ΔmiR(6338)   CGGGUGUCCUGGCUCCAAUUCCACUCAACUGGUAUGUGUUAAAGCCUUGGUGUAGCAGCGGUUGUUACUCUCCAGUGUACUCG
               ((..((((..((((..(((.(((......))).....)))..)))).(((.(((((((...)))))))...))).........)
```

```
ebv-miR-BART7
                          ebv-miR-BART7*                     ebv-miR-BART7
               ((((((..(((.((((((((((.((((((((..((............))..)))))))).)).)))))))).)))..)))).))
AJ507799       CCAGUGUCCUGAUCCUGGACCUUGACUAUGAAACAAUUCUAAAAAAAUGCAUCAUAGUCCAGUGUCCAGGGACAGUGCACUCGG
               ||||||||||||| |  ||      |     ||| ||||||||||||||   |   ||     |   |  ||||||||||||||
r_ΔmiR(6338)   CCAGUGUCCUGAUUCGAGAUUGACGCAUCACAACUAUUCUAAAAAAAUGGCCCUAUGUUAGCCGGAAACUGACAGUGCACUCGG
               .....)))).......((((((..((...(((..(((((.......)))))).....((((((.......))))))))))....)).
```

```
ebv-miR-BART8
                          ebv-miR-BART8                     ebv-miR-BART8*
               (((((.(((((.((((((..((.(((((((((((((..........)))).))))))))).))..)))))).))))).)))))
AJ507799       UGGGUUCACUGAUUACGGUUUCCUAGAUUGUACAGAUGAACUAGAACUGUCACAAUCUAUGGGGUCGUAGACAGUGUGCUUA
               |||||||||||||   |      |  |        ||||||||||||||    | ||| | | |    |  |||||||||||
r_ΔmiR(6338)   UGGGUUCACUGAUCUGGCAGCUUUUUACAAGUUGAAUGAACUAGAACUGAUGUAGUCUUUCGAGGGCAAACCAGUGUGCUUA
               (((((.(((((.(((((((((((.....))))))......)))))).........((((.....))))....))))).)))))
```

Figure S7 continued

```
ebv-miR-BART9
                              ebv-miR-BART9*                      ebv-miR-BART9
               (((.((((((..(((.((((((((((.((...((((..........))))..)).)))).)))))).)))..))))))...)))
AJ507799       AGCUGUUGUUUGUACUGGACCCUGAAUUGGAAACAGUAACUUGGAUUCUGUAACACUUCAUGGGUCCCGUAGUGACAACUAUGCU
               |||||||||||||  ||      |          ||||||||||||||||      || | |       |   ||||||||||||
r_ΔmiR(6338)   AGCUGUUGUUUGUCUUGAGGAUAGCCAACAGCUAAGUAACUUGGAUUCUGCCGACUUUGACGCUGUGAAUUCAGACAACUAUGCU
               .(((((((.(((((.....))))).))))))).......((((......)))).......))..)).(((((.((....))))
```

```
ebv-miR-BART22
                                                                  ebv-miR-BART22
               ((((..(((((((.(((((((((.(((..((((((.........)))).)))..))).))))))))).))))))).))))
AJ507799       GCGGUUGUCACAGGUGCUAGACCCUGGAGUUGAACCAGUACCACUCGGUUACAAAGUCAUGGUCUAGUAGUUGUGACCCUGC
               ||||||||||||||||||||||||||||||||||||||||||||||            |   | | |||  | |  ||||||
r_ΔmiR(6338)   GCGGUUGUCACAGGUGCUAGACCCUGGAGUUGAACCAGUACCACUCAAGCUGUCGUGGAGAUUAUGGUACAUAUUGCCCUGC
               ..((((.((.((((........))))))....)))).(((((((((..((....)).)))....))))))...)))))))...
```

```
ebv-miR-BART10
                              ebv-miR-BART10*                     ebv-miR-BART10
               ((((((..(((.(((((.((((.(((((.((((((..........)))))).))))).)))).))))).)))..))))))
AJ507799       GGAGUGUCCCGGGGCCACCUCUUUGGUUCUGUACAUAUUUUGUUAUUGUACAUAACCAUGGAGUUGGCUGUGGUGCACUCC
               ||||||||||||||||||||||||||||||||||||||||||||||            |   | | |||  | |  ||||||
r_ΔmiR(6338)   GCGGUUGUCACAGGUGCUAGACCCUGGAGUUGAACCAGUACCACUCAAGCUGUCGUGGAGAUUAUGGUACAUAUUGCCCUGC
               ..((((.((.((((........))))))....)))).(((((((((..((....)).)))....))))))...)))))))...
```

```
ebv-miR-BART11
                              ebv-miR-BART11-5p                    ebv-miR-BART11-3p
               ((((.((((((((.(((.((((((((((.((..(((((.......)))))...)).)))))))))).))).))))))))...))))
AJ507799       GGCUUCUGUUGGGUCAGACAGUUUGGUGCGCUAGUUGUGUGCUUAGCAGCAACGCACACCAGGCUGACUGCCUUAGCAGUGUGGCC
               |||||||||||||         |        | | | |||||||||||||||| |        | |  | ||||||||||||||
r_ΔmiR(6338)   GGCUUCUGUUGGGCGUUUGGAGUGUCGUAAAUCGGUCUGUGCUUAGCAGCAACCCGCUGUGCCAAGCCGACAUUAGCAGUGUGGCC
               ((((.((((((((((.......)))).....((((..((.((...(((((.....))))))))..))))...)))))))...))))
```

```
ebv-miR-BART12
                                                                  ebv-miR-BART12
               ((((((.(((((.(((((((...((((((.((((((.........)))))).)))))))...)))).))).))))).))).)))
AJ507799       CUGGUGACCUAACACCCGCCCAUCACCACCGGACAGAUUCUGAACUUGUCCUGUGGUGUUUGGUGUGGUUUUGGGGUACGCAG
               |||||||||||||||||||||||||||||||||||||||||||||||| | | ||  |||  |||| |||||||||||||||
r_ΔmiR(6338)   CUGGUGACCUAACACCCGCCCAUCACCACCGGACAGAUUCUGAACUUGCGCGGGGGGUUUUUUUGUGUUUUUGGGGUACGCAG
               (.((((......)))).))..........((((.....))))....)))))))).((.(((((((((.........))))))
```

Figure S7 continued

```
ebv-miR-BART19
                            ebv-miR-BART19-5p                    ebv-miR-BART19-3p
              (((.(((((((...(((((((.(((((((.((((((............)))))).)))))))..)))))))...))))))).)))
AJ507799      CCGUGUCCUGACAACAUUCCCCGCAAACAUGACAUGGGUUAAUUUAAACAUGUUUUGUUUGCUUGGGAAUGCUCUUAGGGCCUGG
              |||||||||||||     ||    | |||   | | ||||||||||||||||||| ||  | ||  |   |||||||||||||||
r_ΔmiR(6338)  CCGUGUCCUGACACUCGACCAAAGAGACACCUCUUAGGUUAAUUUAAACAUGUUUAGUGCGGUUUUGUUAGCUCUUAGGGCCUGG
              (((.(((((((..........(((((....))))).(((((((..((((.(((.....))))))).))))))).))))))).)))
```

```
ebv-miR-BART20
                         ebv-miR-BART20-5p                 ebv-miR-BART20-3p
              (((((.((.(((((((((.(((((((((.((.............)).)))))))))..))))))))).)).))).)))
AJ507799      AGGGCCUAUUGUAGCAGGCAUGUCUUCAUUCCUGCGUACCGAAUGGCAUGAAGGCACAGCCUGUUACCAUUGGCACCU
              |||||||||||   |          ||   | |||||||||||||| | |  | |    |      | ||||||||||
r_ΔmiR(6338)  AGGGCCUAUUGCUACUCUGUCAAGAUCUGGCUUGCGUACCGAAUGGUAGGCCGCCGGUACAACAAUCUAUUGGCACCU
              ((((((....((((.(((....)))..)))).....((((((..(((....))).))))))...........))).)))
```

```
ebv-miR-BART13
                           ebv-miR-BART13*                      ebv-miR-BART13
              (((((.(((.((.(((.(((.((((...((((((............))))))....))))))).))).)).))).)))))
AJ507799      GGCACCUCGAUAACCGGCUCGUGGCUCGUACAGACGAUUGUUUGGCUCUGUAACUUGCCAGGGACGGCUGACGAUGUGUU
              |||||||||||     |  | |    |    |||||||||||||||||        |       |||   ||||||||||
r_ΔmiR(6338)  GGCACCUCGAUUCGGCGGCCAUUAACCCGGUAGACGAUUGUUUGGCUCGCACGUCAGUUCAUAGCGGGGUACGAUGUGUU
              (((((.(((((((.(((((((......((.....))......)))).)))......((.....)).)))).))).)))))
```

```
ebv-miR-BART14
                             ebv-miR-BART14*                  ebv-miR-BART14
              ((((.(((.((((..(((((((((((..(((((((..........)))))))...))))).))))))..))))...)))..))))
AJ507799      CAGGGGUGGCCGGUACCCUACGCUGCCGAUUUACAUAAUAUAAAUUGUAAAUGCUGCAGUAGUAGGGAUCUGGACGCGCGACCUG
              |||||||||||||                |     ||||||||||||    |  ||  | | |    | ||||||||||||||||
r_ΔmiR(6338)  CAGGGGUGGCCGGCCUAUACUCGCCGAACUGCUAUUAAUAUAAAUUGGGGUUCGUGAUGGAAUCUAAAACUGGACGCGCGACCUG
              (((.((((((((((........))))....))))))..........((((((......))))))....)))))))..((.....)
```

```
ebv-miR-BART2
                            ebv-miR-BART2-5p                     ebv-miR-BART2-3p
              (((((..(((..((((((((.((.(((((((((((..........))))))).))))..)).))))))))..)))..)))))
AJ507799      GGACUUCCAGACUAUUUUCUGCAUUCGCCCUUGCGUGUCCAUUGUUGCAAGGAGCGAUUUGGAGAAAAUAAACUGUGAGUUU
              ||||||||||||     |            || ||||||||||||||||||  |  | | |   |    |||||||||||||
r_ΔmiR(6338)  GGACUUCCAGACCGAACUUCUUUCCGUGUUUUCCGUGUCCAUUGUUGCAAGAUGAAAAUCGAUUAGGGGAAACUGUGAGUUU
              ((((....((((((..........)).)))).....))))..........(((....)))..(((.((....)).)))....
```

Figure S7 continued

| *AJ507799:* | miR-BHRF1-1<br>nt 41464-41544 | miR-BHRF1-2<br>nt 42840-42920 | miR-BHRF1-3<br>nt 42956-43039 |
|---|---|---|---|

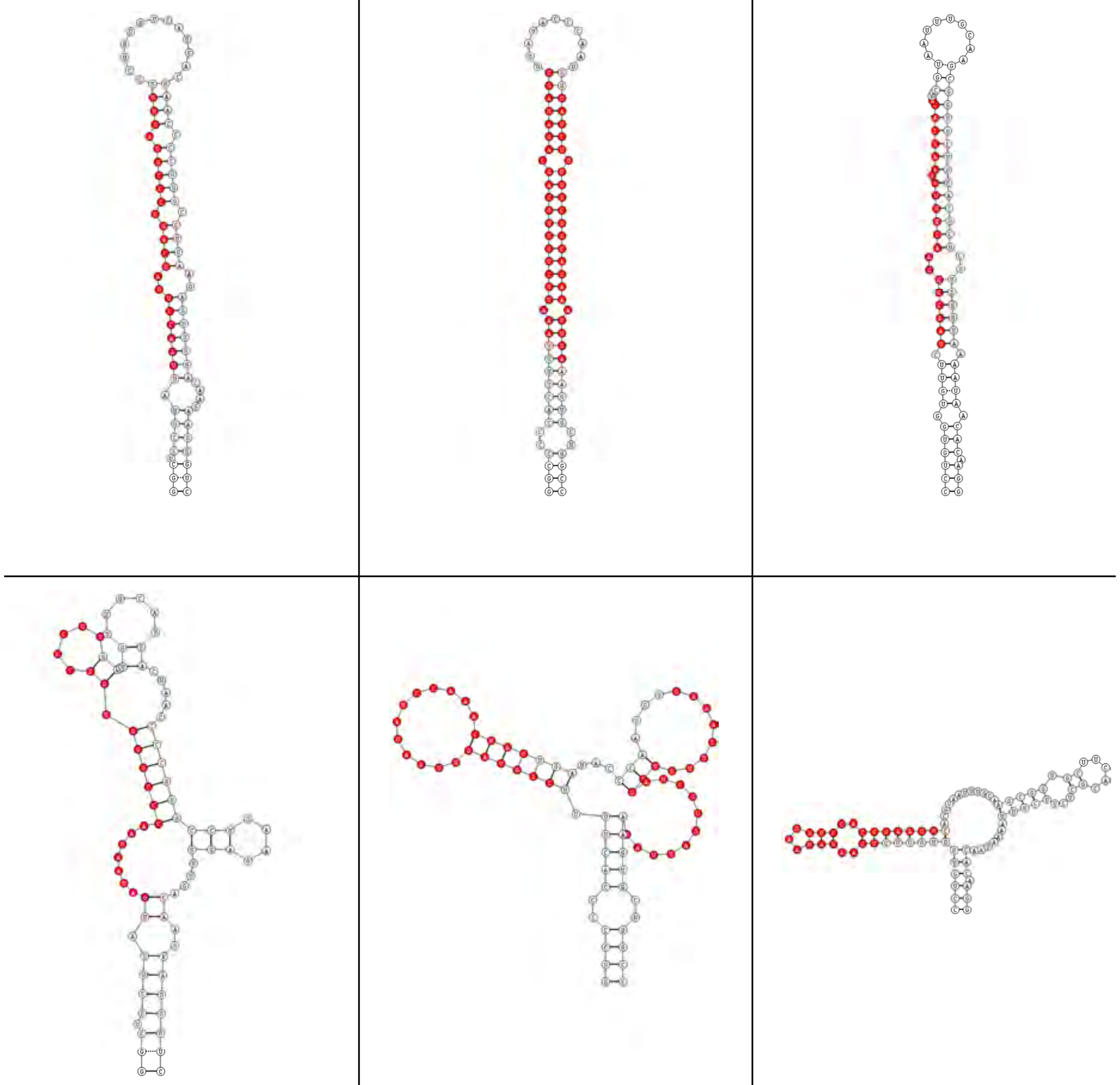

| r_ΔmiR (6338) | r_ΔmiR (6338) | r_ΔmiR (6338) |
|---|---|---|

Figure S7 continued

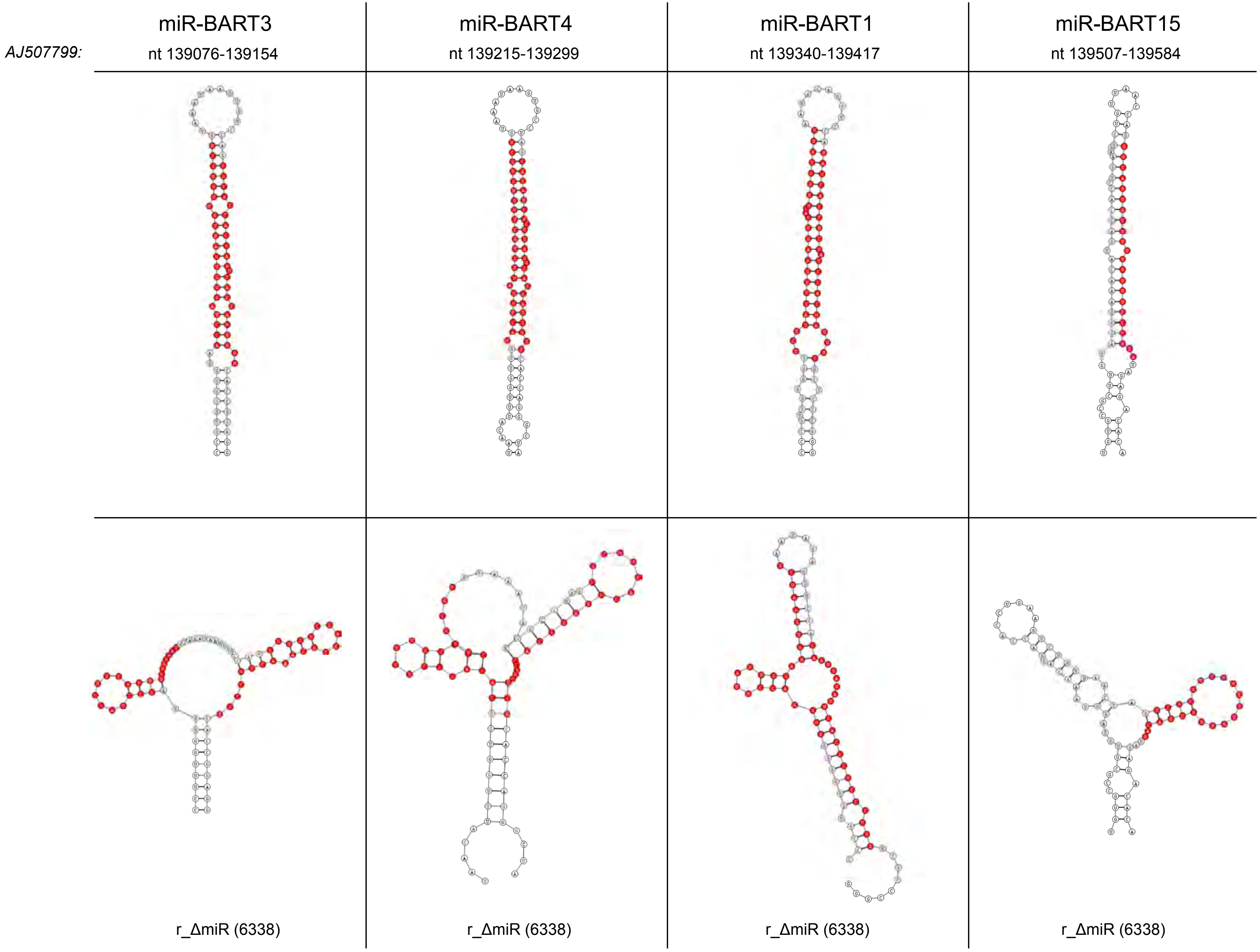




Figure S7 continued

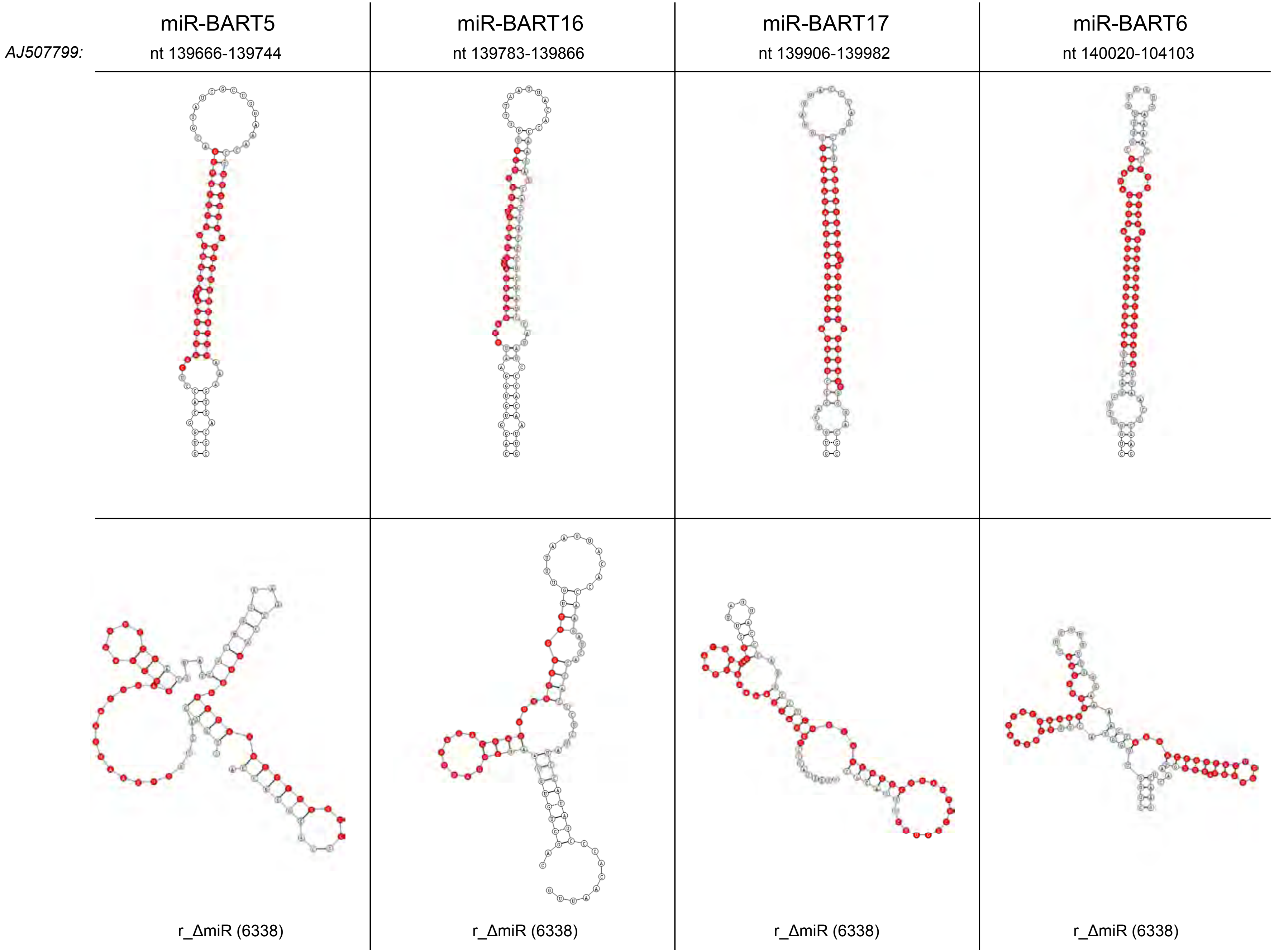




Figure S7 continued

| | miR-BART21 | miR-BART18 | miR-BART7 | miR-BART8 |
|---|---|---|---|---|
| *AJ507799:* | nt 145499-145583 | nt 145949-145583 | nt 146420-146502 | nt 146759-146840 |

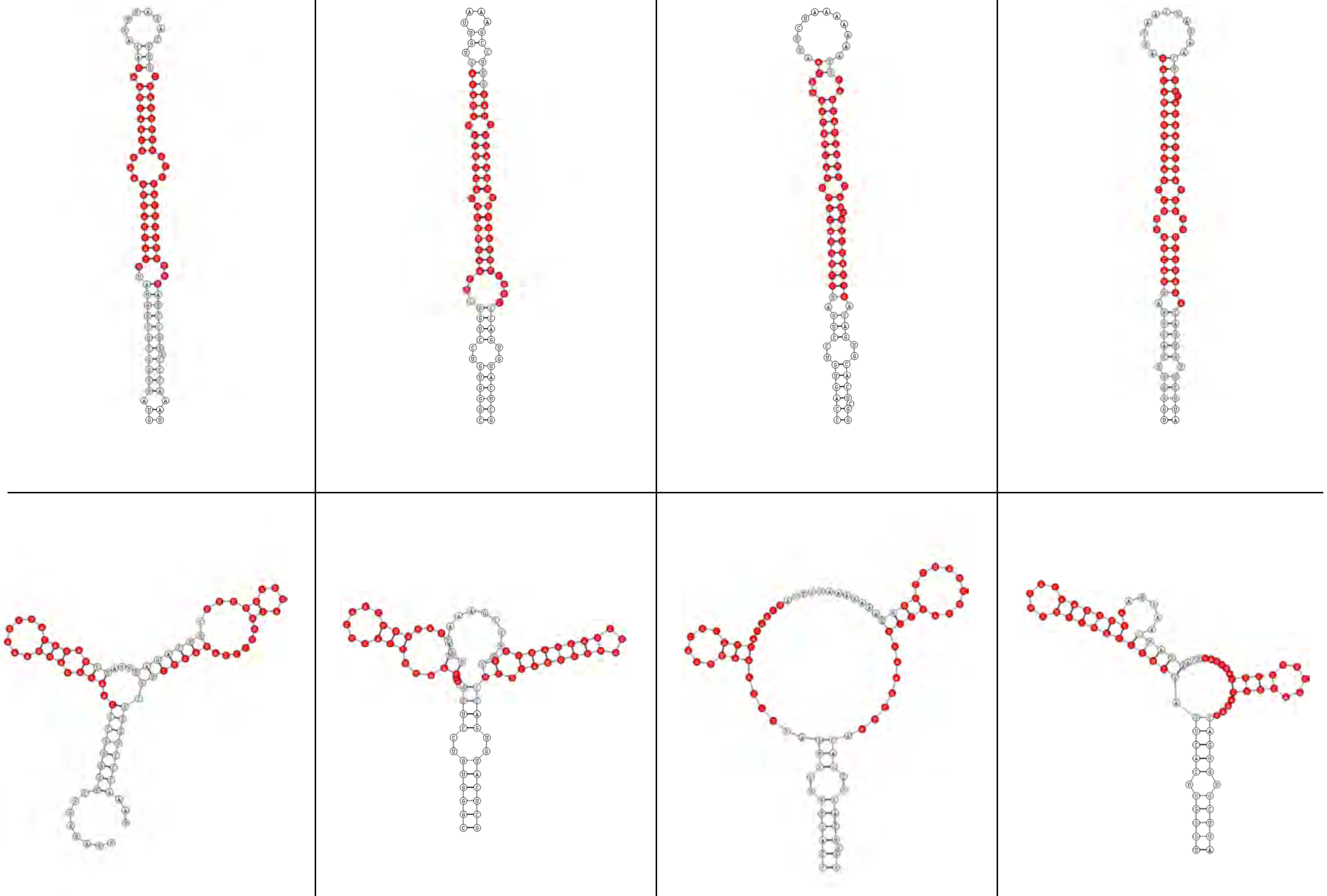

| r_ΔmiR (6338) | r_ΔmiR (6338) | r_ΔmiR (6338) | r_ΔmiR (6338) |
|---|---|---|---|



Figure S7 continued

| | miR-BART9 | miR-BART22 | miR-BART10 | miR-BART11 |
|---|---|---|---|---|
| *AJ507799:* | nt 146947-147031 | nt 147155-147236 | nt 147308-147388 | nt 147524-147609 |
| | 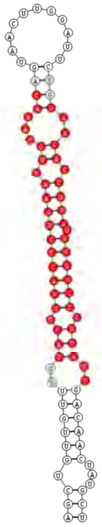 | 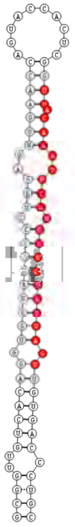 | 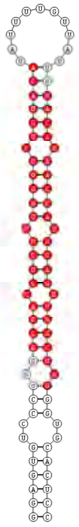 | 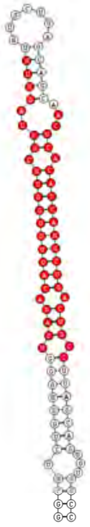 |
| | 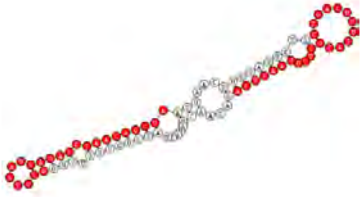 r_ΔmiR (6338) | 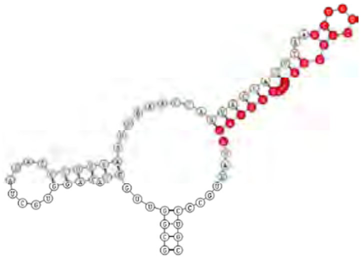 r_ΔmiR (6338) | 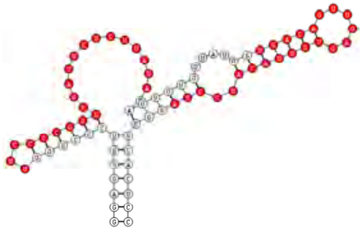 r_ΔmiR (6338) | 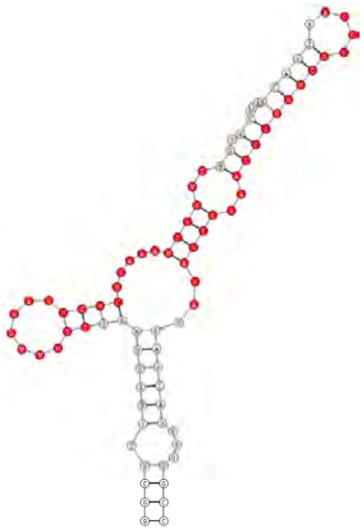 r_ΔmiR (6338) |



Figure S7 continued

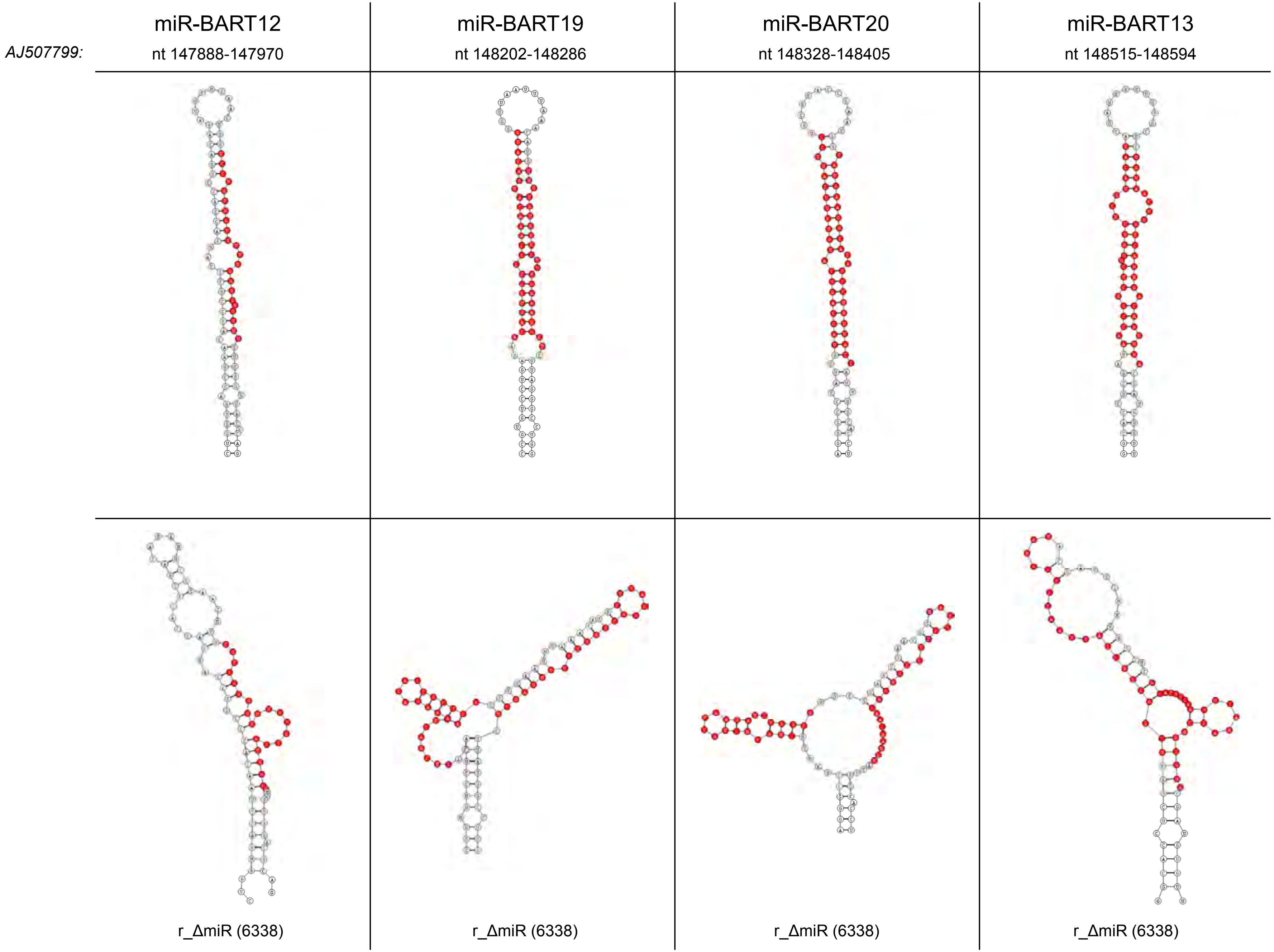




Figure S7 continued

| | miR-BART14 | miR-BART2 |
|---|---|---|
| *AJ507799:* | nt 148731-148815 | nt 152735-152816 |
| | 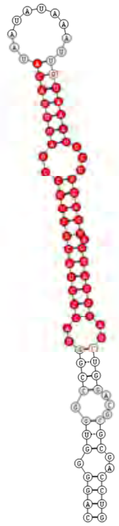 | 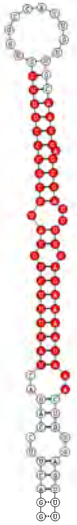 |
| | 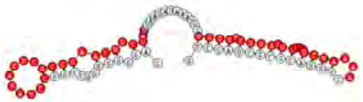 | 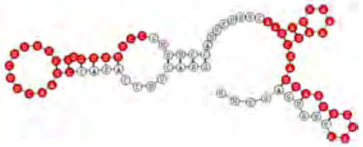 |
| | r_ΔmiR (6338) | r_ΔmiR (6338) |

Figure S7 continued
